# Adaptation to mutational inactivation of an essential E. coli gene converges to an accessible suboptimal fitness peak

**DOI:** 10.1101/552240

**Authors:** João V. Rodrigues, Eugene I. Shakhnovich

**Affiliations:** Department of Chemistry and Chemical Biology, Harvard University, 12 Oxford Street, Cambridge, MA 02138, USA

## Abstract

Genetic inactivation of essential genes creates an evolutionary scenario distinct from escape from drug inhibition, but the mechanisms of microbe adaptations in such cases remain unknown. Here we inactivate *E. coli* dihydrofolate reductase (DHFR) by introducing D27G,N,F chromosomal mutations in a key catalytic residue with subsequent adaptation by serial dilutions. The partial reversal G27->C occurred in three evolutionary trajectories. Conversely, in one trajectory for D27G and in all trajectories for D27F,N strains adapted to grow at very low metabolic supplement (folAmix) concentrations but did not escape entirely from supplement auxotrophy. Major global shifts in metabolome and proteome occurred upon DHFR inactivation, which were partially reversed in adapted strains. Loss of function mutations in two genes, *thyA* and *deoB*, ensured adaptation to low folAmix by rerouting the 2-Deoxy-D-ribose-phosphate metabolism from glycolysis towards synthesis of dTMP. Multiple evolutionary pathways of adaptation to low folAmix converge to highly accessible yet suboptimal fitness peak.

## Introduction

When important cellular functions are inactivated, e.g. by genetic mutations, long ranging disruptions of cellular networks can occur, which poses a major adaptive challenge to cells. Recent studies investigated evolution of *E. coli* upon inactivation of non-essential enzymes of carbon metabolism that lead to re-wiring through less efficient pathways (1–6). These studies highlight a crucial interplay between regulatory responses and imbalances in metabolite concentrations resulting from gene knockouts, which can be subsequently corrected by mutations elsewhere. Importantly all these studies involved gene-knockout so that adaptation by reversal to wild type genotype was not possible.

Less is known about the adaptation mechanisms that follow inactivation of unique cellular processes that are deemed indispensable in microbes. The genes that perform such functions, often classified as essential, are believed to face higher selective pressure and thus evolve slower (7). It is thus expected that disruption of essential genes either by mutation or by antibiotic stress can create a major barrier to the adaptability of bacteria. Nevertheless, a comprehensive study involving knock outs of >1000 genes classified as essential in yeast has shown that a small percentage of mutants could recover viability after laboratory evolution (8), revealing that essentiality can, in some cases, be overcome through adaptation. Loss of essential biosynthetic genes have also been observed to occur naturally, under evolutionary conditions where pathway products are provided externally (9), highlighting a context-dependent nature of essentiality. Nevertheless, while adaptation upon inactivation of essential genes in microbes has been demonstrated, its mechanisms remain unknown.

Here we study evolutionary adaptation upon functional inactivation of an essential *E. coli* enzyme dihydrofolate reductase (DHFR). Past efforts to link fitness effects of chromosomal variation in the *folA* locus encoding DHFR and their biophysical effects on DHFR protein allowed us to develop an accurate quantitative biophysical model of DHFR fitness landscape (10–14). The availability of a clear genotype-phenotype relationship makes DHFR a unique model to study the dynamics and outcomes of evolution to recover from its inactivation by point mutations. In contrast to previous studies that used gene knockouts here we inactivate the chromosomal *E. coli* DHFR by introducing mutations in the *folA* locus at a key catalytic residue in position 27, generating strains that express DHFR inactive protein (with at least 4 orders of magnitude lower catalytic efficiency than the wild type) but are viable only with external metabolic compensation, allowing the mutants to adapt to lack of DHFR function by decreasing supplement concentration. The advantage of this approach is that it presents cells with an obvious evolutionary “ solution” of reverting the mutant back to wild type variant without massive rewiring that could lead to potentially lesser fit variants. However, an actual outcome may depend on evolutionary dynamics which could revert to wild type or other form of active DHFR (higher fitness peak) or converge to a potentially more accessible solution of rewiring to a less efficient metabolic pathways that do not require DHFR function. Therefore, this setup allows us to assess relative roles of height and accessibility of fitness peaks in determining the outcome of evolutionary dynamics.

We show that, depending on starting DHFR variant, partial reversion of DHFR phenotype may indeed occur. However, adaptation to low concentration of external metabolites through metabolic rewiring is the prevalent evolutionary solution due to the availability of a greater number of trajectories leading to consecutive gene inactivation events in two key loci, *thyA* and *deoB*. Using omics analysis, we observe global perturbations in metabolites and proteins levels occurring due to DHFR inactivation and upon adaptation, highlighting a key role of regulatory circuits in directing evolution. Finally, we show how adaptation to loss of drug target generates highly resistant strains, and that one important evolutionary solution found here, inactivation of thyA gene, can be also found in clinical isolates of resistant strains of *staphylococcus aureus* (15, 16) and *haemophilus influenzae* (17).

## Results

DHFR is encoded by the *folA* gene and is essential for the biosynthesis of purines, dTTP, glycine and methionine. We sought to inactivate *folA* gene product in *Escherichia coli* by introducing mutations in the key catalytic residue D27 (Figure 1A).

**Figure 1.**
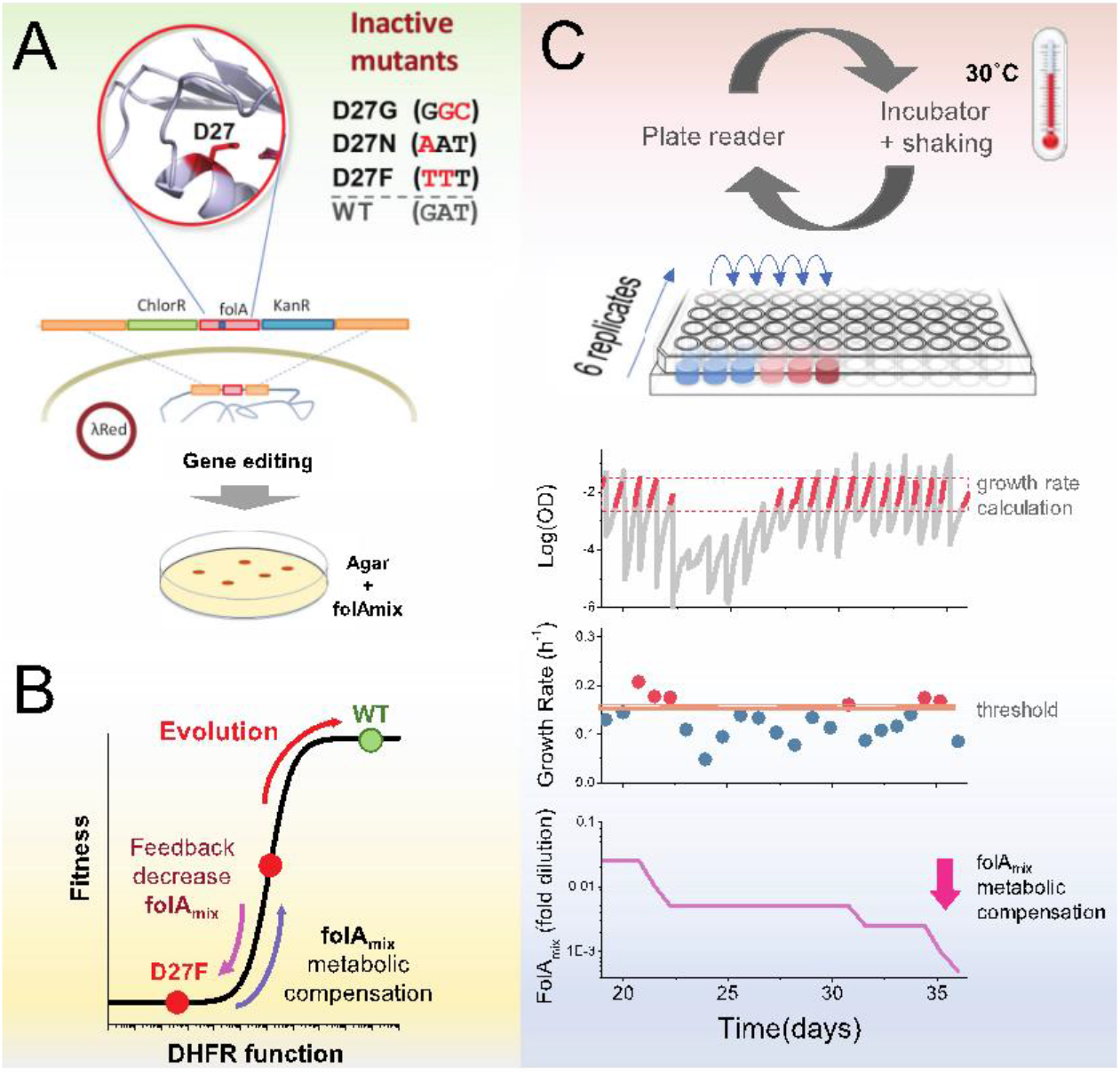
Automated experimental evolution of inactive DHFR mutants. A) Mutations in key D27 catalytic residue of DHFR were introduced in E. coli chromosome, together with flanking antibiotic resistant markers, by lambda red recombination and strains devoid of DHFR function were selected with antibiotics in folAmix-containing agar plates. B) D27 mutants are lethal but fitness can be partially rescued by metabolic compensation with folAmix. Adaptative changes that increase fitness can be counterbalanced by decreasing the concentration of folAmix in the growth medium, forcing the cells to evolve without DHFR function. C) Experimental evolution using automated liquid handling. Six replicates of cell cultures are placed in the first column of a 96well plate and incubated at 30°C with shaking and the optical density is measured every 30 min. The cultures grown until the average OD reaches a threshold (0.3), that was defined to avoid nutrient limitation and consequent shift into the stationary phase. At this point cultures are diluted simultaneously into the wells of the adjacent column, to a common starting OD (0.01), and the cycle is repeated. The growth rate is calculated for every culture and whenever it exceeds a defined threshold the amount of folAmix is independently decreased for each culture in the subsequent dilution.

To construct these strains we supplemented the growth medium with a mixture (folAmix) of adenine, inosine, thymine (precursor of dTTP), methionine and glycine which allows cells to overcome DHFR deficiency (18, 19). We envisaged that inactivated DHFR mutants growing in the presence of supplement folAmix could be progressively challenged to adapt to ever decreasing concentrations of this supplement in the growth medium (Figure 1B). Next, we implemented a fully automated serial passaging scheme as depicted in Figure 1C-D using a Tecan liquid handling robot. In this setup, the growth rate of replicate cultures is monitored by periodic OD readings, and the concentration of folAmix is adjusted downward in each serial passage when cultures exceed a defined growth rate threshold. This feedback control loop ensures mutant strains are continuously challenged to grow at sub-optimal conditions, sustaining a strong selective pressure on the loss of DHFR function. This approach combines the medium-throughput capabilities of plate-based serial dilution methods (currently, up to 48 independent replicate trajectories can be evolved in parallel), and both real-time monitoring of fitness status and control of growth conditions featured in morbidostat setups (20, 21), which allows cells to continuously experience exponential growth and sustained selection pressure.

### D27 mutations confer growth defects

We selected single or double nucleotide base mutations to replace a key catalytic residue Asp 27 either for Asn, Gly or Phe; the presence of a carboxylate side chain in position 27 is strictly conserved among 290 bacterial DHFR orthologues (Figure S1A). Purified D27 mutant proteins were characterized (Table 1)

**Table 1.**
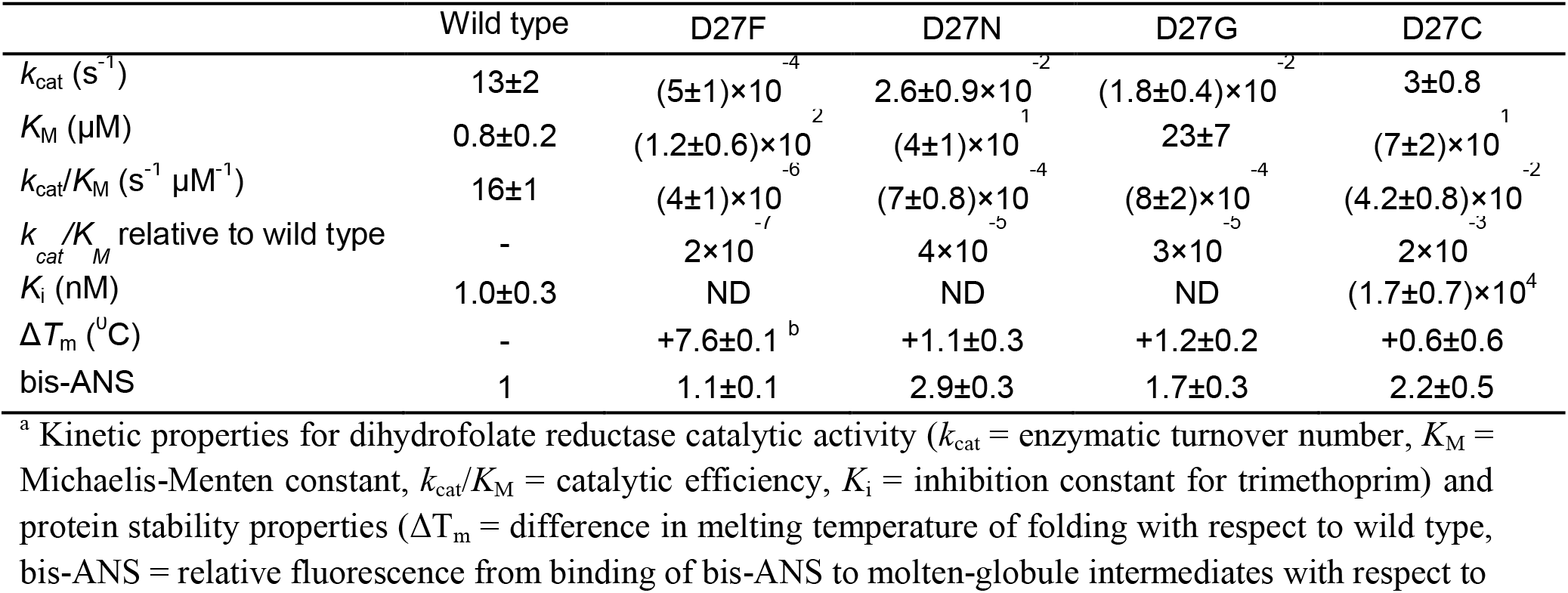
In vitro properties of DHFR mutant proteins

The catalytic activity measurements confirm the lack of significant DHFR function in these variants; catalytic efficiencies (*k***cat**/*K***M**) are several orders of magnitude lower than wild type. However, thermal denaturation data obtained for the D27 mutant proteins indicates that their stability is mostly unaffected (or even significantly increased in the case of D27F mutant (22)) showing that, despite the lack of catalytic activity, these proteins retain the ability to fully fold. Importantly, this provides a pathway for restoration of DHFR function over the course of evolution, either by revertant mutations at the D27 locus or, potentially, compensatory mutations elsewhere in the protein. D27 mutant strains were generated by lambda red recombination (13) and plated in folAmix-containing agar media, however, growth was only visible after 48h and the colonies formed were miniscule. When DHFR function was assayed in cell lysates of D27 mutant strains, no DHFR activity could be detected (figure S1 B,C), confirming that these mutations inactivate DHFR function in the cell. As expected, growth experiments showed that D27 mutants grow only at high concentrations of folAmix (Figure 2A).

**Figure 2.**
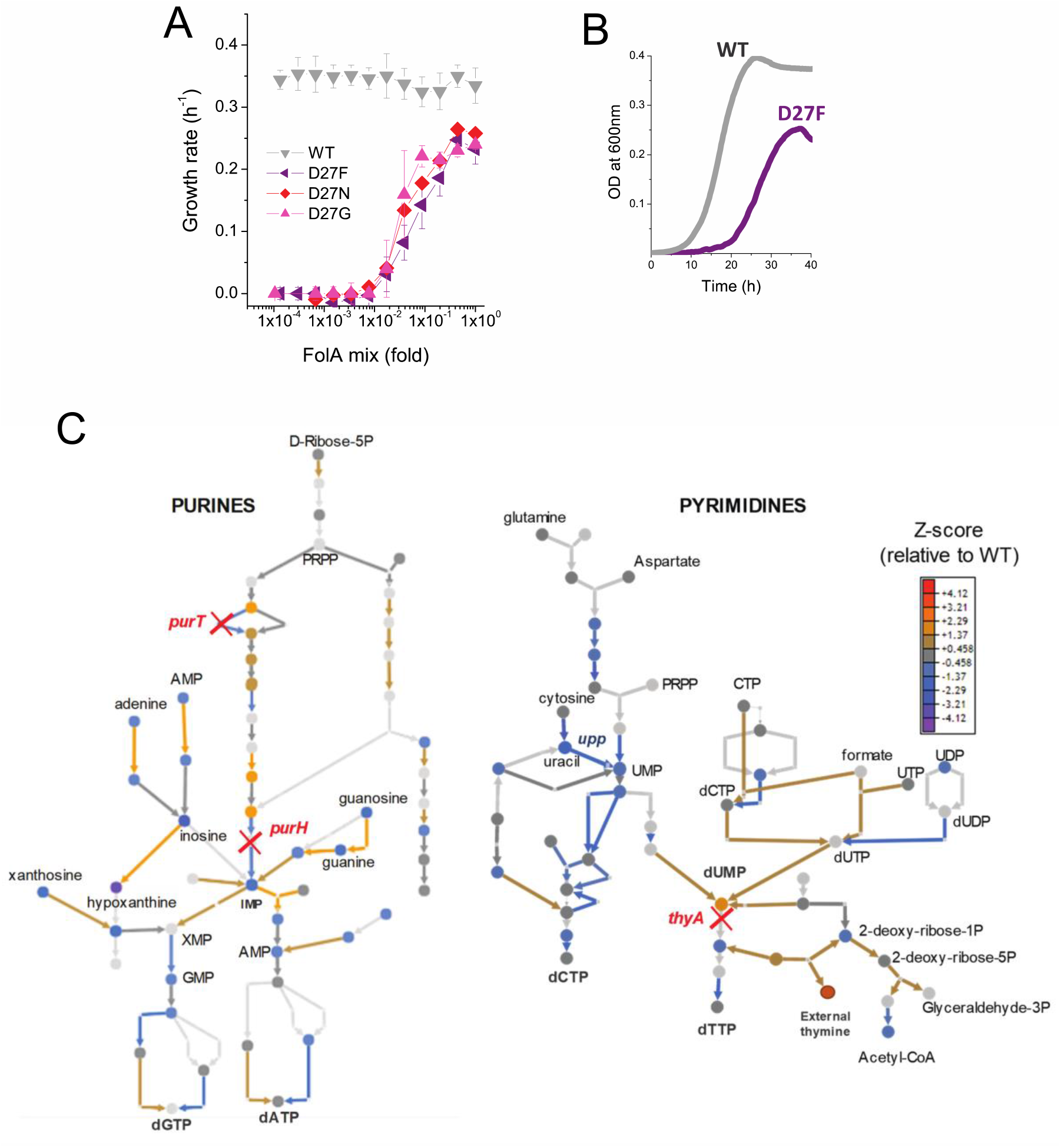
D27 mutants require folAmix to grow. A) The growth rates of each D27 mutant and wild type were measured at various dilutions of folAmix, with respect to initial composition(adenine 20 μg/mL, inosine 80 μg/mL, thymine 200 μg/mL, methionine 20 μg/mL and glycine 20 μg/mL), (see also Figure S1 and S2). Cultures were grown in M9-minimal medium at 30°C, and absorbance was monitored at 600nm. Data are represented as mean ± SD (N=4) B) Comparison of representative growth curves of wild type and D27F mutant obtained in the presence of 1x folAmix. C) Schematic representation of purine and pyrimidine biosynthetic pathways depicting changes in the levels of metabolites and proteins for the naïve D27F mutant (Z-scores, relative to wild type). Metabolites are represented by circles and arrows represent enzymatic reactions color-coded by the levels of associated proteins (see also Figure S3 and S4). Light gray shading indicates that no data is available. Metabolites that are downstream of the enzymatic reactions requiring reduced folates (ThyA, PurH and PurT) are strongly depleted in naïve D27F mutant, whereas those metabolites immediately upstream to those reactions are increased in respect to wild type.

Interestingly, D27N and D27G mutants show slightly better growth at lower concentrations of folAmix than D27F. Given that the catalytic efficiency of D27G and N, albeit extremely low, is 2 orders of magnitude higher than that of D27F, it is possible that such difference could have a beneficial impact at low folAmix concentrations. In addition, potential acquisition of a slightly advantageous mutation elsewhere upon genetic manipulation to introduce D27G/N mutation in the *folA* locus can also lead to growth differences (see Discussion). We tested D27 mutants for growth in the presence of individual components of folAmix and their different combinations and found that only thymine was essential, although growth with thymine alone is slower compared to growth with all folAmix components (Figure S2A,B). The growth rates measured for all D27 mutants at high folAmix fall in the range 70-80% of wild type and lag times were typically 1.5-2 fold longer (Figure 2 B).

### D27 mutations cause major metabolic and proteomic changes

The previous observation that D27 mutant strains show severe growth defects suggests that DHFR inactivation imposes considerable homeostatic imbalance. We focused on the strain D27F to carry out a detailed characterization of the systems-level effects of DHFR inactivation. To that end we carried out high throughput proteomics analysis of D27F mutant strains using LC/MS TMT approach as described earlier (12). The method based on differential labeling provides abundances of proteins in the proteome relative to a reference strain which in our case was wild type (12). We computed Z-scores of the log of relative (to wild type as reference) abundance according to the following equation:

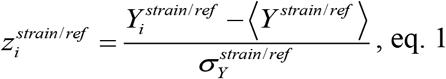

where index *i* refers to protein, 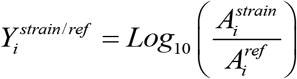 where 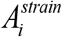 and 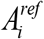 are the protein i abundances obtained for the mutant and reference (wild type) strains, respectively, 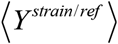 denotes a quantity 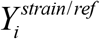 averaged over all proteins for a given strain, and 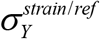 is the standard deviation of ***Y**^strain/ref^* (see Dataset S1). Then, we classified the proteins by their cellular function according to Sangurdekar et al (23) and performed a Student t-test to determine which groups of proteins had statistically significant variation of protein levels in naïve D27F strain, with respect to wild type (see Dataset S2). Numerous cellular processes were significantly altered, reflecting broad genome-wide effects of DHFR inactivation (Figure S3). Proteins involved in central processes were significantly downregulated in naïve D27F strain, including energy metabolism (aerobic respiration, TCA cycle), metabolism of nitrogen, several amino acids, pyrimidines and lipopolysaccharides. On the other end, we found significant increase in the expression of proteins involved of stress responses, peptidoglycan recycling and salvage of guanine and xanthine. We then performed both targeted and untargeted metabolomic analysis of naïve D27F mutant strain and wild type (see experimental details) to characterize significant changes at the level of metabolites. Likewise, Z-scores for metabolite levels were computed for naïve D27F mutant, with respect to wild type (see Dataset S3). The metabolites with the highest absolute Z-scores (>1.96) were selected for an enrichment test using MBRole online software (24) to identify pathways in which altered metabolites are overrepresented. The analysis revealed that the metabolism of purines, pyrimidines, beta-alanine, histidine and sulfur were the most significantly changed in naïve D27F mutant comparatively to wild type (Figure S4A,B). A detailed scheme representing the changes in metabolites and proteins levels of nucleotide synthesis pathways is shown in Figure 2C. Not surprisingly, we observe build-up of metabolites upstream the reactions that require reduced folate cofactors (PurT, PurH, and ThyA), whereas metabolites downstream of those reactions are found to be strongly depleted.

Overall, weaker growth and small colony phenotype show that D27 mutants are severely compromised even in presence of folAmix. Using the evolutionary scheme described previously (Figure 1C), we allowed six replicates derived from the same colony of each mutant strain D27F/N/G to evolve in parallel for about 50 days.

### Evolution of D27G mutant

Cultures of D27G mutant strains were first grown in the presence of folAmix diluted to 0.1 fold with respect to the initially defined folAmix composition.

Throughout the course of the experiment the concentration of supplement mixture further decreased (Figure 3A) in response to increasing fitness of the mutant strains, as imposed by the feed-back control loop discussed previously. Thus, the change in folAmix concentration over time reflects the effect of adaptation. Three trajectories (1,3 and 6) stand out by reaching the ability to grow in the complete absence of folAmix. Trajectories 2 and 5 reached a point where growth dropped below detection limit, and ultimately could not be recovered, whereas trajectory 4 had adapted to grow at low concentrations of folAmix. We hypothesized that mutations in *folA* locus could have restored DHFR catalytic activity in the cultures where reversion of phenotype was observed. Accordingly, Sanger sequencing analysis of this region revealed that in all these three trajectories (1,3 and 6), but not trajectory 4, residue Gly27 had mutated to cysteine. This finding was surprising because alignment of multiple known DHFR sequences shows that cysteine does not occur naturally in position 27; only aspartate or glutamate are observed in this locus (Figure S1A). To assess whether Cys27 mutant is functional, this variant was purified and characterized in vitro, and its catalytic properties are compared with wild type and other mutants in Table 1. Although stability-wise D27C mutant is similar to wild type protein, it shows much weaker catalytic efficiency, mostly in terms of *K_M_* retaining only about 0.1% *k*_cat_/*K*_M_ of wild type. To assess if this residual catalytic function is sufficient to explain the reversion of phenotype in the evolved strain we first predicted fitness based on a previously established model (10). Taking as input the vitro biophysical properties of purified D27C mutant, namely kcat/KM and bis-ANS fluorescence values, the model predicts a growth rate of about 30% of the wild type strain. To check the validity of this prediction, the mutation D27C was reconstituted in the wild type background by lambda red recombination and cells were plated in folAmix-containing agar plates to lift any selective pressure for DHFR activity. This strain was able to grow in the absence of folAmix supplementation, and the measured growth rate was 92% of the wild type, which is in fair agreement with the in vitro-based prediction which does not take into account a possibility of upregulation in response to DHFR deficiency (12) (see SI for discussion) (Figure 3B-D). This analysis shows that D27C mutation alone explains significant fraction of fitness recovery of the evolved strains. Of note is also the fact that significant loss in catalytic efficiency in D27C mutant pays-off with a dramatic 4 orders of magnitude increase in the inhibition constant for trimethoprim, and consequent increase of IC50 for trimethoprim inhibition obtained for D27C strain (Figure 3E). This is in line with previous observations that active site mutations compromise catalytic function but also disrupt pocket interactions with the drug, resulting in high levels of resistance (10, 25).

**Figure 3.**
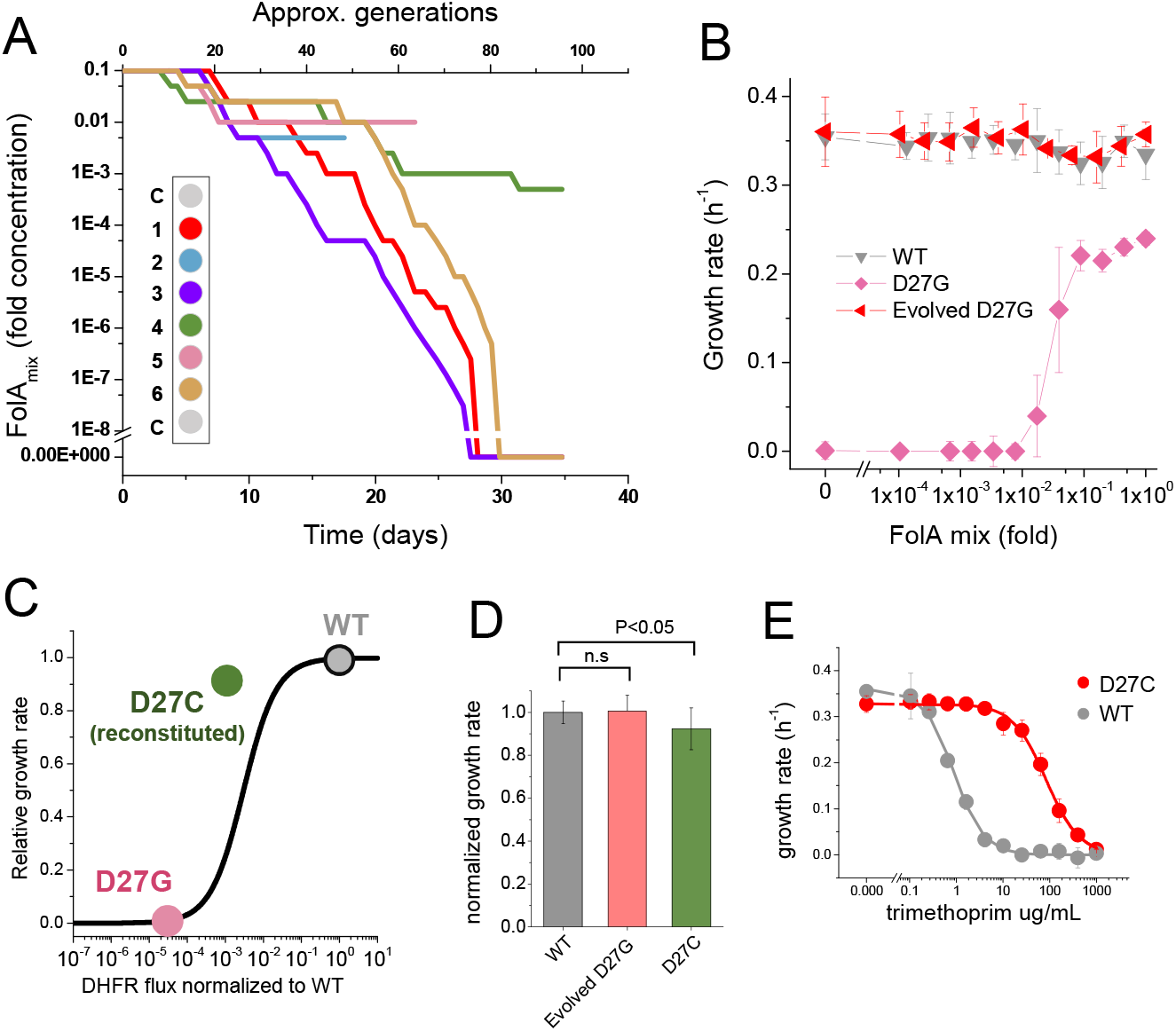
Phenotypereverting G27C mutation emerges upon evolution of D27G strain. A) Evolutionary profiles of the six trajectories in D27G adaptation showing the time dependent changes in folAmix concentration necessary to sustain growth. Each trajectory is colored according to its position in the column of the 96-well plate, as represented in the scheme (C = wells with growth medium only). B) Evolved D27G strain does not require folAmix to grow. The growth rates determined at different concentrations of folAmix are compared for wild type, naïve and evolved D27G strain from trajectory 6 (clone G33T6#1). Data are represented as mean ± SD (N=4). C) Biophysical properties determined in vitro for the D27C DHFR were used to predict growth rate based on the DHFR fitness (10). The resulting prediction yields a growth rate that is 30% of the wild type value. This is in fair agreement with the experimental value measured for a BW25113 strain in which D27C mutation was introduced in the wild type folA gene (D27C reconstituted) which was 92% of the wild type. D) Comparison of growth rates of wild type, evolved D27G and reconstituted D27C strain (mean ± SD, N=4). E) Dose-dependent growth inhibition by trimethoprim determined for wild type and reconstituted D27C mutant (mean ± SD, N=3). Solid lines are fits with logistic equation, from which IC50 was determined to be 0.8 ± 0.1 μg/mL for wild type and 79 ± 17 μg/mL for D27C mutant.

### Evolution of D27F and D27N mutants

Evolutionary trajectories of D27F and D27N mutants represented in Figure 4A and 4B, respectively, show a marked decrease in folAmix necessary to sustain growth, in most cases reaching concentrations nearly three orders of magnitude lower than initially required for naïve strains.

**Figure 4–.**
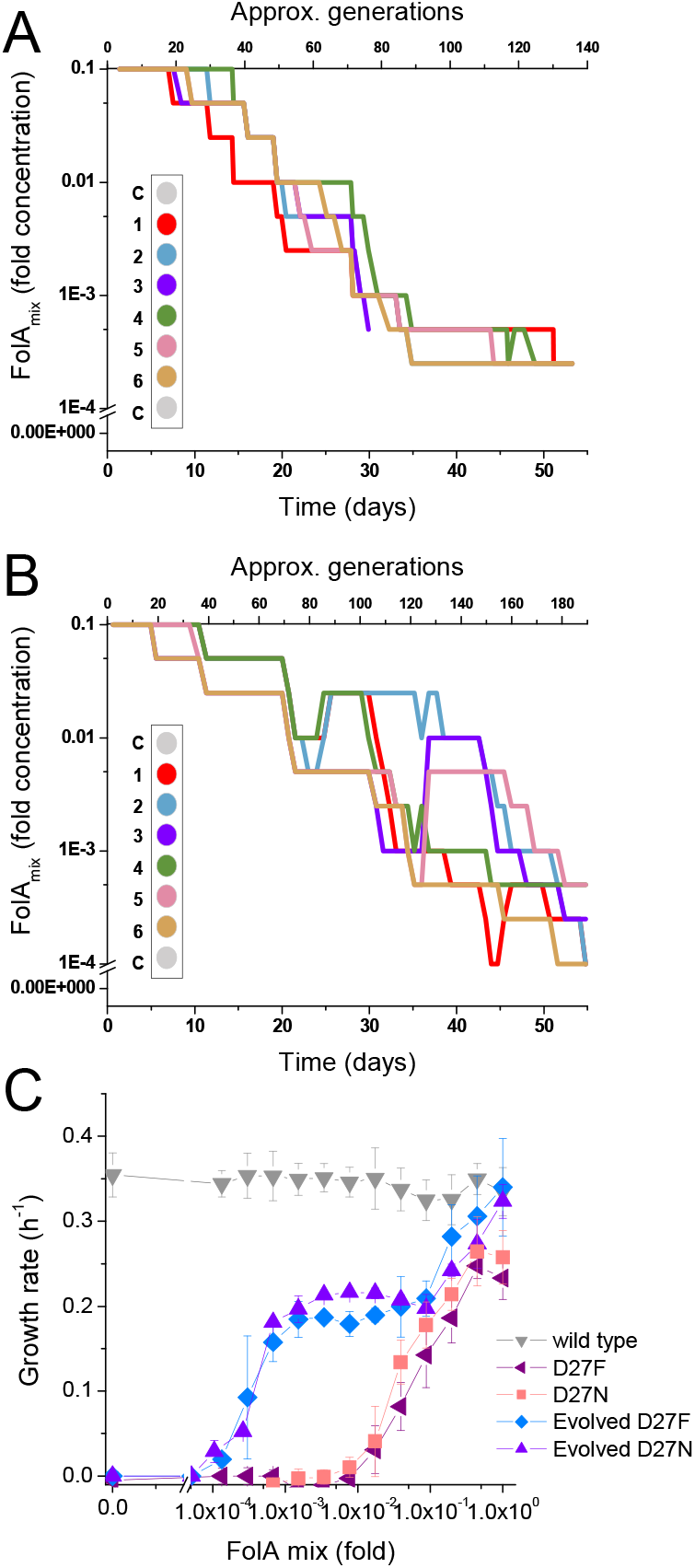
D27F and D27N mutant evolved to grow at low folAmix concentrations. Evolutionary profiles of each trajectory in A) D27F and B) D27N adaptation to loss of DHFR function. C) Comparison of growth dependence on folAmix concentration determined for wild type, both naïve and evolved D27F (trajectory 1, clone F51T1#1) strains and naïve and evolved D27N (trajectory 6, cloneN51T6#1), see also Figures S2, S5. Data are represented as mean ± SD (N=4).

To verify that these strains adapted to grow at low folAmix concentrations we first plated evolved cultures (from −80°C glycerol stocks) in agar medium supplemented with 1x concentration of folAmix and then randomly selected individual colonies from different trajectories to measure growth at various concentrations of folAmix. Plating the cultures at a high folAmix concentration ensures that there is no biased selection for high-fitness phenotypes when colonies are randomly picked from agar media. We found that individual colonies taken among all trajectories that adapted to low folAmix concentrations all showed a strikingly similar phenotype - folAmix-dependent growth profile, irrespective of the initial variant at D27 locus (F/N and also trajectory 4 in D27G) (Figure S2 C-E). Figure 4C shows representative growth profiles obtained with colonies taken from trajectories 1 and 6 of D27F and D27N, respectively, showing a marked decrease in growth rate at concentrations of folAmix comparable to the lowest concentrations achieved during the evolution experiment. However, these strains cannot grow in the complete absence of folAmix. This result indicates that, unlike some trajectories in D27G evolution, adaptation of D27N and D27F strains did not involve reversion of the *folA*^−^ phenotype. Accordingly, no mutations were found in *folA* locus in evolved D27N and D27F strains (see Dataset S4). Other mechanisms must therefore be responsible for the ability to grow at low folAmix concentrations. The growth rate of the evolved strains measured at high concentrations of folAmix reached values comparable to wild type, showing a clear fitness increase with respect to naïve strain. However, at lower concentrations of folAmix the growth rate of evolved strain becomes markedly reduced to nearly half of the wild type value and appears to plateau in an intermediate range of folAmix concentrations. The growth rate value at this plateau coincides with that measured for cells grown in thymine alone (Figure S2A,B) indicative that thymine is the growth-limiting component at these concentrations of folAmix. We also evaluated sensitivity to trimethoprim for both naïve and evolved D27F strains. Not surprisingly, in the absence of a functional DHFR and in the presence of folAmix D27F mutant strains are extremely resistant to trimethoprim (Figure S5A). We noted, however, that while no decrease in growth was evident in naïve D27F strain up to 1000 μg/mL, the growth rate of the evolved strain dropped abruptly above 500 μg/mL trimethoprim, suggesting that the drug is acting on an unknown target in the cell which is essential in the evolved but not the parent strain. We then tested how the resistance of evolved D27F strain would compare with wild type at very low concentrations folAmix (Figure S5B). We found that in those conditions IC50 for the wild type is similar to that measured in the absence of supplement, yet the evolved D27F strain still shows an extremely high resistance to trimethoprim (IC50=656±78 μg/mL).

### Loss of function mutations in thyA and deoB lead to adaptation to low folAmix concentrations

Whole genome sequence (WGS) analysis was performed to identify the genetic basis for the adaptation mechanisms of the evolved strains. We analyzed naïve D27F/N/G strains and individual clones representative of adaptation to low folAmix (evolved D27F - trajectories 1 and 2 and evolved D27N - trajectory 6), and one clone representative of adaptation through DHFR reversion (D27G- trajectory 6). In addition, to characterize the dynamics of mutation fixation, individual colonies from intermediate evolutionary time points of D27F trajectory 2 were also sequenced. Figure 5 A-B summarizes the WGS results obtained for each clone of evolved D27 and naïve strains.

**Figure 5–.**
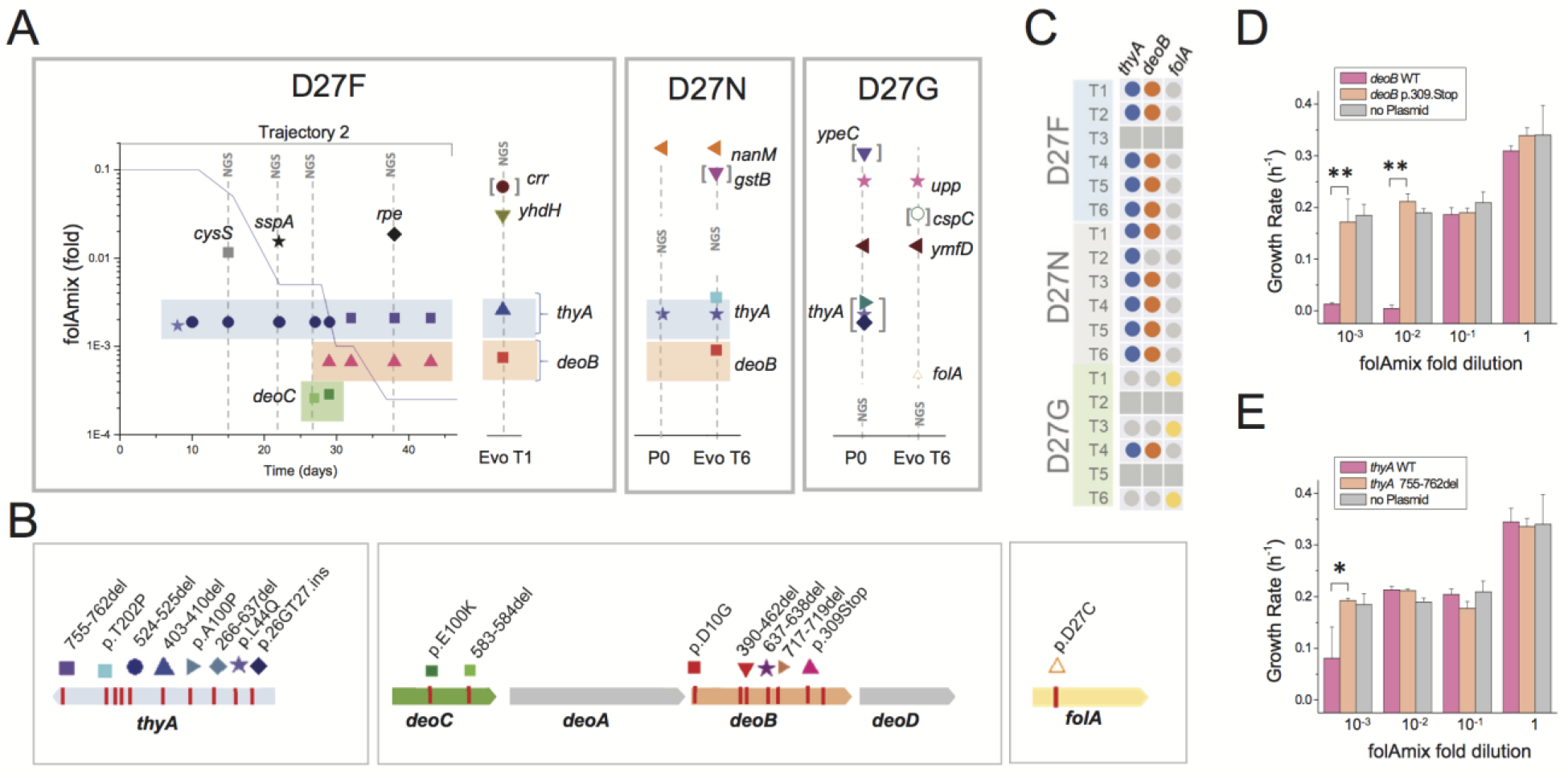
Loss of function mutations in two genes, thyA and deoB, lead to adaptation to low folAmix concentrations. A) Mutations identified in naïve and evolved D27 mutant strains. Vertical lines identified with NGS represent mutations identified by whole genome sequencing whereas the remaining cases were identified by Sanger sequencing. Details are presented for trajectory 2 of D27F evolution and include the mutations identified at various passages and the folAmix profile obtained for that trajectory. Points denoted P0 correspond to naïve strains. Mutations between brackets represent polymorphisms with estimated frequencies of 0.5. See also Figure S6 for the effect of upp and ymfD mutation in naïve D27G strain. Figure B) Mutations identified in the most relevant genes. C) Mapping thyA, deoB and folA mutations found among evolved strains from all trajectories of each D27 mutant. Colored dots represent the presence of mutations, whereas gray dots represent genes that were not mutated in evolved strains. Trajectories that ceased growth prior to the end of the experiment were not sequenced (gray squares). D-E) Expression of wild type DeoB and ThyA in evolved D27F strain reverts phenotype to high folAmix requirement. Evolved D27F strain (trajectory 1, clone F51T1#1), with background mutations deoB p.D10G and thyA 403-410del, was transformed with pTRC-tetR plasmid coding D) wild type or mutant deoB and E) wild type or mutant thyA, under the control of tetR repressor. Evolved D27F strain without plasmid is also represented as control. Cells were grown in M9 minimal medium and growth rates were measured from periodic OD measurements.. * P<0.01, ** P<0.001.

We found that D27N and D27G strains, but not D27F, had additional background mutations that must have been acquired prior to starting the evolution experiments; although all D27 strains were constructed from the same parent wild type strain, we could not prevent the appearance of mutations at any given stage prior to the evolution experiments. It is possible that these mutations may provide some fitness advantage with respect to “ pure” D27F strain (see below and discussion). Comparison between evolved and naïve D27G strains reveals that no other mutation was fixed upon evolution besides the one nucleotide change at the D27 locus of the *folA* gene, corresponding to the Gly->Cys substitution described earlier. This is fully consistent with the earlier conclusion that D27C mutation alone explains the fitness recovery observed in D27G evolution. We asked if the presence of mutations in *upp* and *ymfD* genes originally found in D27G strain could have an impact on fitness of the cells, and thus could have conditioned the evolutionary fate of this strain. We note the potential role of the mutation in *upp* gene, which encodes the enzyme uracil phosphoribosyltransferase that participates in pyrimidine salvage pathway and, quite strikingly, appeared depleted in proteomics analysis of naïve D27F strain. To address this, we engineered additional D27G strains and, after confirming the absence of mutations in *thyA*, *upp* and *ymfD*, compared their growth properties at various folAmix concentrations with D27G *upp/ymfD* strain that was used in the evolution experiments. No difference was found in the growth rate profiles of these strains (Figure S6), indicating no obvious impact of mutations in *upp* and *ymfD* in favoring the reversion of DHFR function over *thyA* inactivation.

From the analysis of evolved D27F and D27N we can identify two common hotspots for mutations, the genes *thyA* and *deoB*, encoding thymidylate synthase and phosphopentomutase, which are involved in the synthesis of dTMP via de novo and salvage pathways, respectively. For that reason, the sequence of these two loci were determined by Sanger analysis for all other trajectories that evolved to grow at low folAmix concentrations, including trajectory 4 in D27G evolution. Strikingly, in a total of 12 trajectories all were found to have mutations in *thyA* and 11 had *deoB* mutated (Figure 5C). We noted as well that evolved D27F strains also carried mutations in other genes, *crr* (7bp deletion), *rpe* and *yhdH* (4bp deletion), however, these were not observed among other trajectories indicating that these are most likely either random passenger mutations or provide only marginal advantage on a specific genetic background. On the other hand, the occurrence of deletions and deleterious point mutations (see Table S1) in both *thyA* and *deoB* strongly implies that functional inactivation of these gene products arises as a main mechanism of adaptation to the lack of DHFR function. We reasoned that complementation with either thymidylate synthase or phosphopentomutase should create a significant fitness disadvantage to the *evolved* cells, which can be reverted if these enzymes become inactivated. To verify this prediction, we transformed evolved D27F strains with plasmids expressing either wild type ThyA or DeoB proteins and compared the ensuing growth rates of these strains with transformants carrying the same plasmids expressing functionally inactive ThyA and DeoB, respectively. Expression of functional ThyA and especially DeoB were found to be highly deleterious in *evolved* D27F strains at low folAmix concentrations, as shown Figure 5 D-E, whereas no significant change in growth was observed upon expressing inactive mutants.

Overall, these results show two competing adaptation mechanisms to the loss of DHFR activity, either involving a reverting mutation at the D27 locus or, more prevalently, consecutive inactivation of two genes.

### Loss-of-function mutations in deoB prevent draining of deoxyribose pools into energy production

The gene *deoB* is part of an operon responsible for deoxyribose degradation, that includes *deoA*, *deoC*, and *deoD*, which is under the control of 2-deoxyribose-5-phosphate-inducible *deoR* repressor (26). DeoB catalyzes the interconversion of 2-Deoxy-D-ribose-1-phosphate and 2-Deoxy-D-ribose-5-phosphate, and, while the former is necessary for synthesis of dTMP from thymine by DeoA, the latter can be further degraded in glycolysis, through conversion into acetaldehyde and D-glyceraldehyde 3-phosphate by DeoC (Figure 6A).

**Figure 6–.**
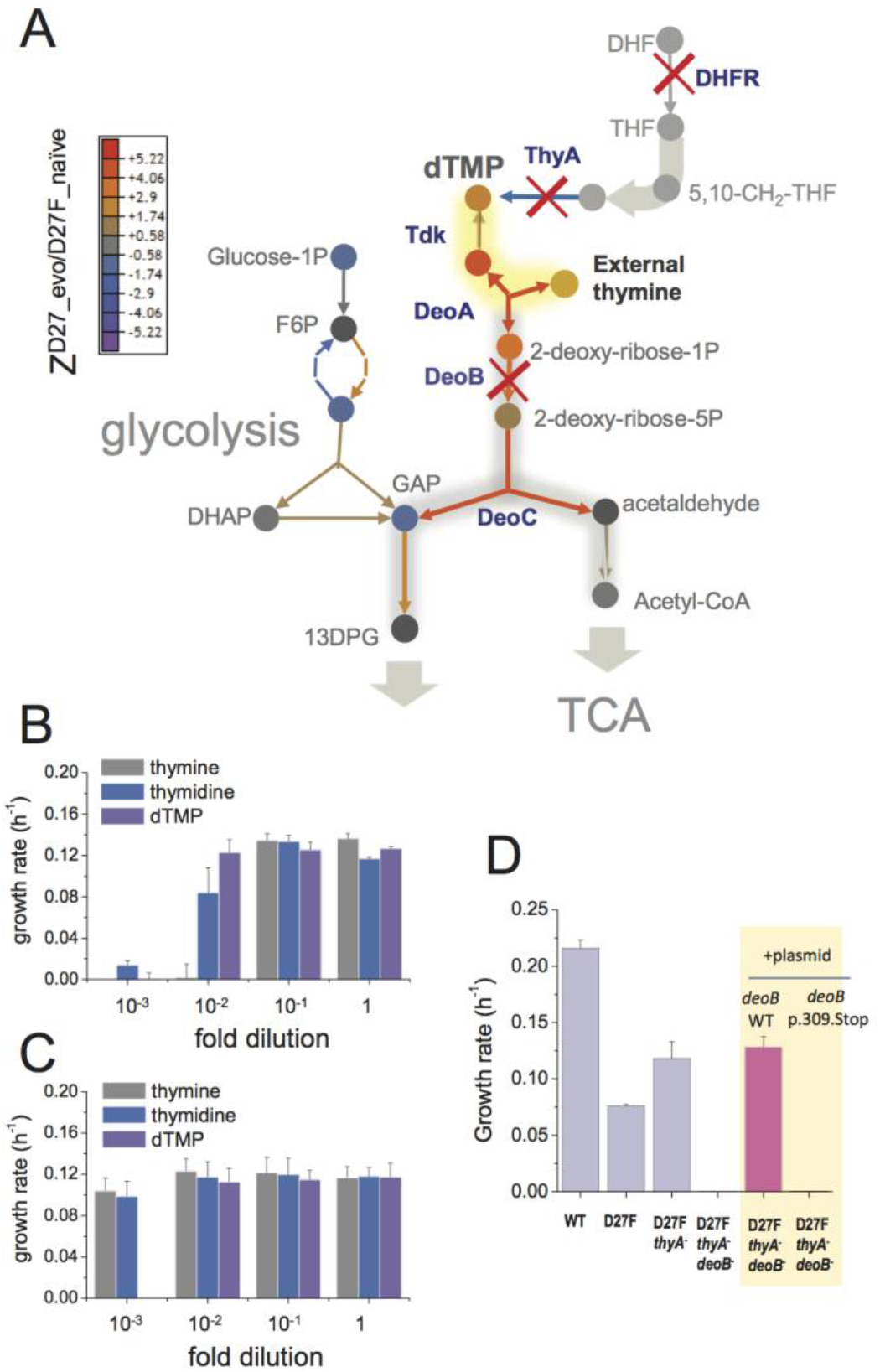
Inactivation in deoB gene prevents 2-deoxy-D-ribose-1-phosphate from being diverted into energy production via glycolysis and tricarboxylic acid cycle. A) Changes in the levels of metabolites (depicted as circles) and proteins (depicted as arrows) in the evolved D27F strain (Z-scores, relative to naïve D27F mutant) are represented. The proteins involved in 2-deoxy-D-ribose-1-phosphate degradation are marginally increased in evolved D27F, with respect to naïve strain (average Z-score = 0.445, P=0.0817, see Figure S7). However, deoB inactivation saves 2-deoxy-D-ribose-1-phospahte that is required in dTMP production, instead of being utilized for energy production in glycolysis, which is significantly upregulated in evolved D27F, with respect to naïve D27F (average Z-score= 0.593, P= 0.015). B) Evolved D27F transformed with a plasmid expressing wild type DeoB are able to growth at low external concentrations of thymidine and dTMP, but not at low concentrations of thymine. C) Evolved D27F strains transformed a plasmid expressing inactive DeoB mutant grow at the same extent with either thymine, thymidine or dTMP. At the lowest concentration of dTMP, however, these cells are not able to grow, likely because of inefficient transport of this metabolite into the cell. D) DeoB^−^ strains cannot grow on thymidine as the sole carbon source unless. Data are represented as mean ± SD (N=3)

Therefore, inactivating *deoB* or *deoC* is an “ economy” solution preventing key metabolite in thymine uptake to be wasted in energy production, allowing cells to efficiently use small amounts of thymine available from media supplement for nucleotide synthesis (27). In agreement with the key role of deoxyribose in limiting thymine uptake, while *deoB*^+^ cells cannot grow at low concentrations of thymine in the media, replacing this precursor by thymidine or dTMP, which provide a deoxyribose moiety, improves the growth of evolved D27F strains transformed with plasmid expressing active DeoB (Figures 6B-C). Moreover, to confirm that the route for deoxyribose degradation into energy production is blocked in evolved mutants, we measured the growth of naïve and evolved cells in the presence of thymidine as the sole carbon source and verified that *deoB*^−^ cells cannot grow unless complemented with a plasmid expressing wild type DeoB (Figure 6D).

### Evolved D27 strains partially revert the omics effects of DHFR inactivation

Next, we carried out LC/MS TMT proteomics analysis of evolved strains (D27F, D27N and D27G) to help identify systems-level changes emerging from adaptation to lack of DHFR function. Since we observed that all clones randomly selected from trajectories that adapted to low folAmix concentrations showed very similar fitness profiles, we focused on individual clones arbitrarily chosen among D27F and D27N trajectories as representatives of adaptation to low folAmix. Specifically, clone F51T1#1 from D27F trajectory 1 and clone N51T6#1 from D27N trajectory 6 were selected. Likewise, clone G33T6#1 from D27G trajectory 6 was chosen as representative of adaptation through reversion of DHFR function. To get a glimpse of global proteomics changes we applied PCA analysis which revealed that both wild type and evolved D27G occupy the same quadrant in the space of two principal components, whereas D27 mutants that are folAmix dependent cluster more closely in another quadrant (Figure S7A). We then computed the Z-scores for the proteomic changes in evolved strains with respect to naïve D27F (Z^D27_evo/D27F_naïve^), to assess the effect of evolution on the proteomic levels and plotted these values against Z^D27F_naïve/WT^ to compare with the initial changes caused by D27F mutation (see also Datasets S1,S2). A clear anticorrelation was observed for all mutants (Figure 7A), being the strongest for evolved D27G strain.

**Figure 7–.**
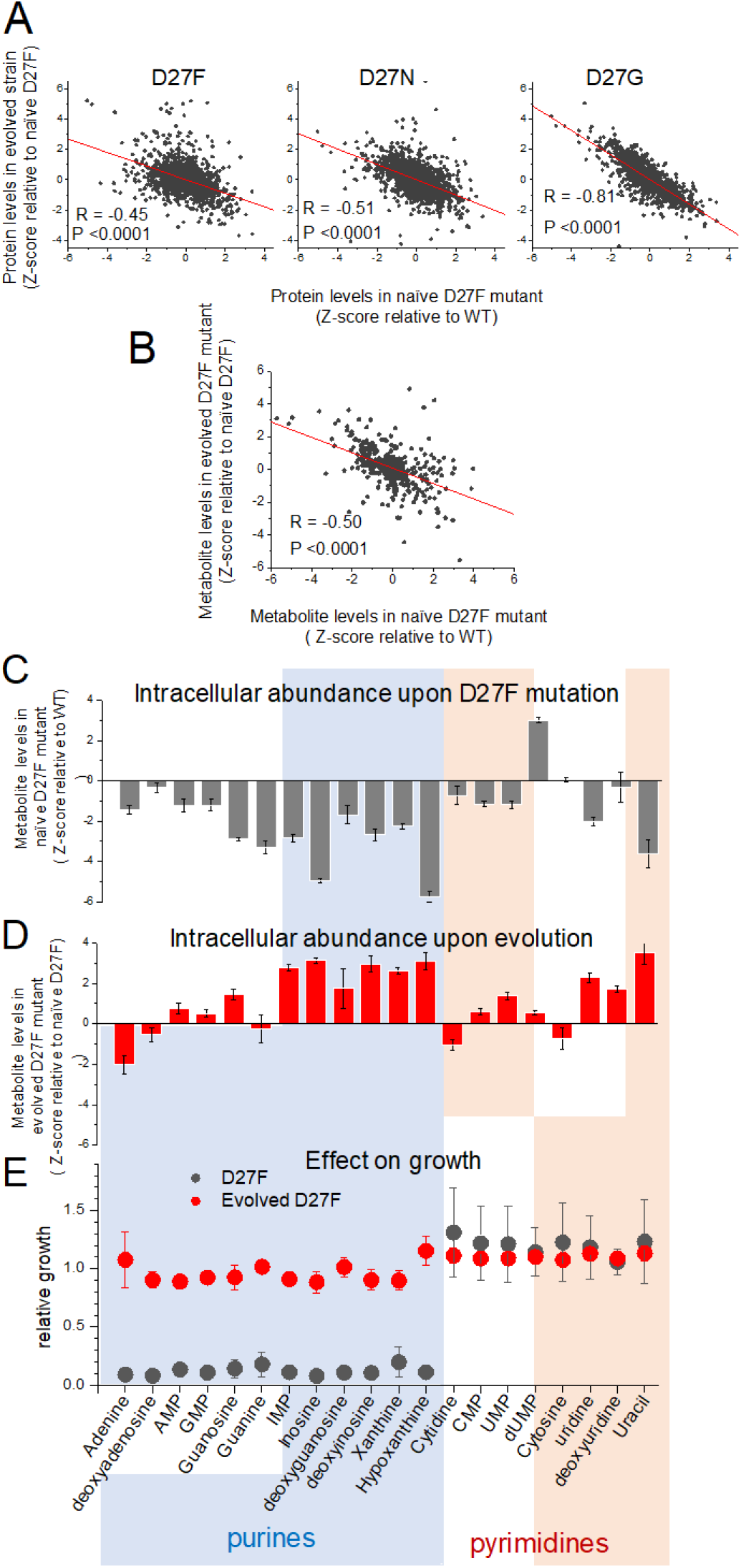
D27 mutation causes major metabolic and proteomic changes. A) Proteomics changes upon evolution partially revert the effect of DHFR inactivation (see also Figures S7 and S4). Comparison of the changes in protein levels obtained for evolved D27F (trajectory 1, cloneF51T1#1), D27N (trajectory 6, clone N51T6#1) and D27G (trajectory 6, clone G33T6#1) (Z-scores, relative to naïve D27F) with those measured for naïve D27F (Z-scores, relative to wild type). B) Comparison of the changes in metabolite levels measured for evolved D27F (trajectory 1, cloneF51T1#1) (Z-scores, relative to naïve D27F) with those measured for naïve D27F (Z-scores, relative with wild type). C-E) Inhibitory effect of purine metabolites supplementation on the growth of the naïve D27F strain is attenuated in the evolved D27F strain. C) Changes in intracellular levels of various purine and pyrimidine species determined by targeted metabolomics for naïve D27F strain, represented as Z-scores relative to wild type. B) Metabolite levels changes obtained for evolved D27F strain, represented as Z-scores relative to naïve D27F strain. C) Effect of individual metabolites on growth of naïve and evolved D27F mutant strains. The effect of supplementation with individual metabolites on growth was determined by growth measurements of naïve and evolved D27F mutant strains in M9 minimal medium supplemented by 1.6 mM thymine in combination with one of each tested metabolites (1 mM final concentration). Relative growth represents the maximum OD obtained for each metabolite normalized to the value measured in the presence of thymine alone. Although most purine metabolites are strongly depleted in naïve D27F mutant, supplementation of the culture medium with these compounds has a strong inhibitory effect on growth. Contrastingly, pyrimidines do not affect growth. In evolved D27F strain, the supplementation with purines has only a marginal effect on growth. Data are represented as mean ± SD (N=3).

These results indicate that adaptation leads to proteomic changes that generally oppose the immediate effects caused by DHFR inactivation. In the case of D27G, the strong global proteomic shift towards wild type levels is somewhat expected since in the evolved strain, the DHFR catalytic activity is partly restored by G27->C mutation. To better characterize the proteomic changes that were specific to evolution of folAmix dependence we searched for classes of proteins grouped by function with significantly altered protein abundance levels in both evolved D27F and D27N with respect to naïve D27F strains (Figure S7B), as described previously. We found a significant increase in the expression levels of proteins involved in glycolysis, fermentation, sugar alcohol degradation and response to low pH, suggesting a metabolic shift towards mixed acid fermentation as a result of adaptation that blocked using derivative of thymidine for glycolysis to save them for DNA synthesis. On the other hand, the levels of proteins involved in DNA restriction/methylation and D-ribose uptake were significantly decreased with respect to naïve D27F in both evolved D27F and D27N strains. Next, we focused on evolved D27F strain to perform a detailed metabolomic analysis and characterize the changes occurring at the level of metabolites (see also Dataset S3). We computed Z-scores for metabolite changes with respect to naïve D27F (Z^D27_evo/D27F_naïve^) and we found significant anticorrelation with Z-scores obtained for naïve D27F (Z^D27F_naïve/WT^) (Figure 7B), indicating that metabolite concentrations in evolved strain generally change to partially recover wild type levels, as observed with proteomics. We found that the pathways significantly enriched in metabolites with the highest Z^D27_evo/D27F_naïve^ scores (>1.96) in evolved D27F were the same as those that had the most significant changes in naïve D27F, with respect to wild type (Figures S4A,B). Overall these results show that adaptation to low folAmix concentrations is accompanied by an overall shift in the abundance of metabolites and proteins to partially revert the system-level changes that are caused by DHFR inactivation.

### Regulatory responses are altered in evolved D27F strain

Metabolites of purine and pyrimidine biosynthetic pathways were among the most strongly depleted in naïve D27F strain and showed highest increase upon evolution (Figures 7C,D and Figure S4A,B). We reasoned that these depleted metabolites could be limiting growth of the D27 mutants, and that supplementing the culture medium with these metabolites would improve growth. Surprisingly, supplementation with purines strongly *inhibited* growth of the naïve D27F strain, whereas pyrimidines addition had, at most, a slight stimulatory effect (Figure 7E). The detrimental effect of individual purine supplementation appears to be associated with a regulatory response to high levels of these metabolites on the culture medium. In the case of evolved D27F strain, the effect of supplementation of individual nucleotides on growth was comparably weaker or non-existent, indicating that the inhibitory effects were relaxed upon adaptation. It is possible that the profound metabolic and proteomic changes upon DHFR inactivation amount to cells switching their proteomes and metabolomes to a novel stable state.

In these conditions, the “ programmed” responses that are triggered by high external concentrations of metabolites seem to actually worsen the fitness of the naïve strain, instead of providing the beneficial advantage that they have evolved for. On the other hand, after evolution, even though metabolites and proteins levels are still quite distinct from wild type values, the regulatory network appears to be better adapted to that new metabolic state. Since we found no mutations directly associated with regulatory genes, it seems that the observed changes result solely from the re-wiring of metabolic pathways. In this regard row examples are particularly noteworthy. Evolution of D27F and D27N strains blocked supplementary glycolysis pathway by rerouting D-oxyribose towards DNA synthesis while the ensuing metabolic changes lead to significant increase in abundance of glycolysis proteins, presumably to compensate for diminished flux of sugar metabolites along the glycolysis pathway (see Dataset S2). Another example of metabolite-induced rewiring is purR operon whereby purR is inhibited by purine hypoxanthine. Evolved strains show significant increase in abundance of hypoxanthine giving rise to significant drop in abundance of proteins controlled by purR (Dataset S2).

Overall, these results show that two key loss of function mutations that are responsible for the partial adaptation of D27F to loss of DHFR activity result in significant metabolic, proteomic and regulatory shifts observed in the evolved strains.

## Discussion

In this study we set to explore the evolutionary mechanisms that follow inactivation of the essential gene that encodes DHFR. Such scenario is akin to antibiotic-mediated target inhibition, but distinct in that there is no selection for mutations that affect interactions with the drug, such as target modification and drug efflux mechanisms. For example, when treated with DHFR inhibitor trimethoprim, bacterial cells recurrently acquire resistance by high-fitness mutations near the active site of the target protein that decrease drug binding affinity, even at the expense of catalytic efficiency (10, 20, 28). In contrast, a genetic perturbation in the same target gene provides the opportunity to study other potential unexplored mechanisms of adaptation in the absence of drug and evaluate their impact in the context of antibiotic resistance. We found that, in the conditions studied here, evolution is constrained towards two solutions that appear to be mutually exclusive, either (partial) restoration of DHFR catalytic activity or adaptation to lack of DHFR function. While D27F, D27N and one trajectory in D27G convergently adapted to lack of DHFR function, other trajectories in D27G mutant reached reversion from thymine auxothroph phenotype. It is important to address the possible reasons for the observed outcome. A potentially influential factor could be the presence of background mutations observed in both D27N and D27G. The point mutation L44Q in *thyA,* reported to affect the activity of a nearly identical orthologue enzyme (29), could have skewed the solution towards further inactivation of *thyA* by subsequent point mutations in other loci of that gene that were later observed in different trajectories. On the other hand, mutations in the genes *upp* and *ymfD* found in D27G strains did not impact the growth rate, thus the role of these mutations in altering the accessibility of the DHFR functional reversion G27->C seems to be neutral. The codon structure of each mutant studied here and bias in the frequency of each mutation type can also affect the likelihood that phenotype-reverting mutations may arise. We note that the nearest accessible high-fitness Asp/Glu codon for D27F mutant is two nucleotide substitutions away (although one nucleotide away from one D27C codon), whereas both D27G and D27N could be rescued by a single G->A or A->G transition, respectively. Only D27G was found to revert, however, not to wild type codon, but to a less catalytically efficient cysteine residue caused by a G->T transversion. In this respect, it is rather surprising that D27C mutation was fixed in 3 trajectories when transversions are generally regarded to be less likely than transitions. That this mutant was able to fix, despite its poor enzymatic activity, brings important insights on how deleterious mutations can be maintained when the selection pressure on protein function is alleviated by particular environmental conditions, in this case the presence of folAmix, and may provide a rapid route to the development of resistance. In addition, these results may hint at a possible mechanism for the divergence of the key residues in the active sites of enzymes across long evolutionary timescales, where fixation of non-consensus mutations, such as D27C, may be allowed by transient adaptation to permissive environments, and subsequent compensatory mutations that restore protein function at later stages.

The most crucial factor in determining the fate of D27 mutant evolution appears to be the higher accessibility of mutational routes for inactivating *thyA* that lead to their fixation very early in the evolution experiment. Thymidylate synthase is the only known enzyme in *E*. *coli* able to synthetize dTMP de novo from dUMP, and inactivation of *thyA* inevitably commits the cell to thymine auxothrophy; reversion of DHFR function on the *thyA*^−^ background is not expected to change this phenotype, as it is known that *folA*^+^ *thyA*^−^ strains are thymine-dependent (30, 31). Competition between DHFR reversion and *thyA* inactivation thus appears to be decisive in determining the evolutionary solution. However, since inactivation of a gene can be achieved by multiple ways, there is greater number of accessible evolutionary trajectories leading to *thyA* knockout than to reversion of DHFR function. This view has major implications for our understanding of how the relative impact of height and accessibility of fitness peaks shapes the evolutionary dynamics short-term adaptation.

Upon *thyA* inactivation, the only available route for dTMP synthesis involves DeoA-catalyzed formation of thymidine from thymine and 2-deoxy-D-ribose-1-phosphate (Figure 6F). However, DeoA is part of the *deo* operon that is induced by high levels of 2-deoxy-D-ribose-5-phosphate. This regulatory scheme allows cells to recycle excess metabolites into energy production. However, on the background of *thyA* inactivation, the co-expression of *deo* operon proteins creates simultaneously a solution and a problem for the uptake of thymine. Now DeoA becomes an essential protein, but its function is opposed by the roles of DeoC and DeoB which divert 2-Deoxy-D-ribose-1-phosphate into degradation via the glycolysis pathway. The evolutionary solution found in D27 strains converged to the inactivation of a single gene of the operon, *deoB*, an event that is sufficient to block the degradation pathway without disrupting the regulatory structure. This critical example illustrates how evolution can be constrained by regulatory circuits that provide conflicting responses when the optimal directionality of metabolic routes is switched due to genetic perturbations. Understanding these processes is critical for guidance towards new strategies for antibiotic resistance prevention. Perhaps not surprisingly, we found that pathways of adaptation to genetic inactivation of an antibiotic target provide a route to high levels of resistance. Likewise, a recent study also reported that *E. coli* cells challenged with increasing concentrations of trimethoprim in the presence of thymidine repeatedly evolved high levels of resistance by *thyA* loss-of-function mutations besides acquisition of commonly observed resistant mutations in DHFR (31). Strikingly, mutations in *thyA* are also known to confer trimethoprim resistance in bacterial clinical isolates of *S. aureus* (15, 16),and *H. influenza* (17), which emphasizes the need of laboratory studies of microbial adaptation as useful models to tackle serious global threat.

More generally this study highlights the evolutionary conundrum between “ survival of the fittest” and “ survival of the fastest”. Unlike previous studies that followed evolution of adaptation to gene knockouts, the current setup provides an “ easy” highest fitness solution by genetic reversion of single amino acid substitution. However, for most evolutionary trajectories the dynamics quickly converged to a “ quick and dirty” solution – a dead-end local fitness peak which provided an immediate relief from stress but blocked the path to highest fitness peak. Future studies will reveal how general this scenario is for evolutionary dynamics of adaptation to mutational inactivation and/or antibiotic induced inhibition of essential bacterial enzymes and beyond.

## Materials and Methods

### Materials

A list of relevant materials is shown in table S2.

### Construction of D27 mutants

Inactive D27 mutants were created by lambda-red recombination (32) according to a previously described procedure (13) with some modifications. Briefly, a pKD13 plasmid was modified to contain the entire regulatory and coding sequence of folA gene, flanked by two different antibiotic markers (genes encoding kanamycin (kanR) and chloramphenicol (cmR) resistances) and approximately 1kb homologous region of both upstream and downstream chromosomal genes flanking folA gene (kefC and apaH, respectively). The entire cassete was amplified using primers PCRseq_KefC_for2 and PCRseq_apaH_rev and transformed into BW25113 cells with induced Red helper plasmid pKD46, and cells were recovered in SOC medium containing 1x folAmix (adenine 20 ug/mL, inosine 80 ug/mL, thymine 200 ug/mL, methionine 20 ug/mL and glycine 20 ug/mL). Transformants were plated in agar media containing both antibiotics and folAmix. To confirm correct integration of the desired mutations, folA locus was amplified by PCR (primers Ampl_RRfolA_for, Ampl_RRfolA_rev) and Sanger-sequenced using primer PCRseq_RRfolA_rev. Plasmid pKD46 was removed by plating cells at 37 °C twice in the absence of antibiotic selection. A list of relevant primers is shown in table S3

### Automated experimental evolution

A liquid handling instrument Tecan Freedom Evo 150 equipped with Tecan Infinite M200 Pro plate reader and Liconic shaker was used in this work. The experiments were done at 30 °C and using M9 minimal media supplemented with 2g/L glucose, 34μg/mL chloramphenicol and 50μg/mL kanamycin. The liquid handling worktable has two 100mL troughs (Tecan) with either fresh M9 medium + antibiotics or 40% glycerol. In addition, solutions of folAmix at various concentrations (prepared in M9 + antibiotics) are provided in 50 mL falcon tubes. The general procedure involves up to four 96-well plates that are used for serial dilutions of bacterial cultures. Each working plate can carry 8 trajectories that are positioned in a single column. In this work we used the first and eight wells in the column with media alone to control for contamination. The experiment starts by placing the 200 μL cultures/control in the first column of the 96-well plate. All the four plates are incubated in a shaker at 30 °C and at every 30 min the OD of each plate is determined alternately. The growth rate of each culture is calculated from OD measurements over time. To ensure proper comparison, the growth rate was computed only from OD values within a specified range (0.1-0.25). When the average OD of the six experimental replicates in a plate exceeds a threshold of 0.30 each culture is diluted into the next adjacent column by mixing a calculated volume of both culture and fresh medium and folAmix so that the initial OD is 0.01 in a total of 200 μL. At this point, the remaining portion of the previous culture is mixed with glycerol in an auxiliary plate and subsequently frozen at −80C. The cycle is repeated throughout the entire experiment. At every passage, the growth rate is compared with a threshold value (0.16 h^−1^) and whenever a culture exceeds this value the concentration of folAmix is halved. The volume of folAmix added in each passage is 10, 5 or 2.5μL.

### Media and Growth Conditions

Cell cultures were recovered by inoculating fresh M9 media medium supplemented with 0.2% glucose and 1x folAmix with a portion of −80°C glycerol stocks taken at different passages. After recovery for several hours, the cultures were plated in agar media containing 1x folAmix, 34μg/mL chloramphenicol and 50μg/mL kanamycin and incubated at 30°C. Plating evolved cultures at high folAmix concentrations ensures that the clones growing in agar media are not biased towards high fitness phenotypes. Individual colonies were randomly picked, grown in M9-media+folA mix and stored in glycerol at −80°C for later analysis. Clones selected in this manner were labeled using the following mask: XdTt#*i*, where X refers to the amino acid letter in D27 locus (F/N/G), d is day at which the culture was passaged in the evolution experiment, t is the trajectory number (1–6) and *i* is the unique number of different colonies taken from the same culture. Cell cultures grown overnight were diluted in fresh M9 minimal media containing 1x folAmix and antibiotics and were grown for additional 4-6 h. Cultures were then pelleted by centrifugation and washed 3 times in fresh M9 without folAmix. Microplates containing 150 μL of M9 minimal with 0.8g/L glucose, antibiotics, and varying concentrations of folAmix were then inoculated with each culture at a starting OD of 0.0005. Growth measurements were performed in a Infinite^®^ 200 PRO plate reader for 48h at 30°C with constant shaking. Growth rate values are represented as mean ± SD from at least 3 biological replicates.

### Effect of DeoB and ThyA expression

The genes *deoB* and *thyA* and corresponding mutants *deoB* p.309Stop and *thyA* 755-762del were cloned into a modified pTRC plasmid in which the *laqI* and promoter region were replaced by the *tetR* repressor gene and promotor region derived from plasmid Plasmid pEM-Cas9HF1-recA56 (addgene Addgene plasmid # 89962 (33)). Transformed strain cells were grown in M9 minimal media + folAmix, then washed with fresh M9 media without folAmix and plated at different concentration of folAmix. Growth rate values are the mean ± SD of at least 3 biological replicates.

### Metabolite extraction and LC-MS analysis

Cultures of mutant strains (naïve and evolved D27F) and wild type (30 mL) were grown in parallel in M9 minimal media supplemented with 2 g/L glucose in a 250 mL flask at 30C. At an OD of 0.2-0.25 the cultures were centrifuged. The volume of culture to be pelleted was calculated so that the product OD × volume of cell culture (mL) = 5. The pellet was mixed with 300 μl of 80:20 ratio of methanol:water that had been pre-chilled on dry ice. Samples were vortexed and incubated in dry ice for 10 min followed by centrifugation at 4°C for 10 min at maximum speed. The supernatant was collected, and the pellet was processed by repeating the procedure. Samples were stored at −80°C until analyzed by mass spectrometry. At least three independent biological replicates were analyzed for each strain. LC-MS analysis in the positive and negative mode was performed as previously described (34). Retention times of several metabolites of the nucleotide biosynthesis pathway were determined from the analysis of pure compounds.

### LC/MS TMT Proteomics

Cell cultures from isolated colonies were grown at 30°C in M9 minimal media supplemented with 0.2% glucose and were collected by centrifugation during mid-exponential phase (OD ~ 0.2-0.25), well below saturation due to oxygen limitation (typically at OD>0.8-0.9). Global proteomic analysis was performed as described previously (12).

### Whole-genome sequencing

Sequencing was performed on isolated colonies on Illumina MiSeq in 2×150 bp paired-end configuration (Genewiz, Inc., South Plainfield, NJ). The raw data were processed with the DRAGEN pipeline (35)(35)(35)(34) on default settings, using the E. coli K-12 substr. MG1655 reference genome (GenBank accession no. NC_000913.3). After the alignment completed, structural variations (SV) including large INDELs (above 50) were detected using the program Manta(36).

### Metabolomics data analysis

Data analysis was performed using the software packages MzMatch (37) and IDEOM (38) for untargeted analysis. For peak assignment in untargeted analysis, IDEOM includes both peak m/z values and predicted retention times calculated based on chemical descriptors (39). A list of 32 experimentally measured retention times was initially used to calibrate retention time predictions. The retention time of putatively identified metabolites were found to correlate fairly well with the values included in IDEOM (R^2^=0.73) and published in other studies (40) (40) (R^2^=0.88 and R^2^=0.61, respectively). For this reason, additional metabolites from those sources that closely matched IDEOM assignments were treated as standards in the identification routine. Following identification, Z-scores with respect to wild type (Z^D27F_naïve/WT^) or to naïve D27F strain (Z^D27F_evo/D27F_ naïve^) were calculated using equation 1. Z-scores determined for each set *i* of *n* biological replicates were combined for each metabolite *met* using the expression 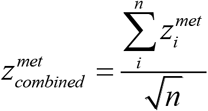. The set of metabolites with highest absolute combined Z-scores (>1.96) was selected to perform an overrepresentation (enrichment) analysis of categorical annotations using MBrole2.0 (24), using as reference the metabolic pathways of *E. coli* from KEGG database.

### Proteomics data analysis

Z-scores were computed using equation 1. For grouping analysis, we used the functional and regulatory classification group sets as (23). For each set of genes belonging to a group we employed a one-sample t-test which provides the p-value against the null hypothesis that the group of genes was drawn from a normal distribution and considered that a given group of genes is upregulated or downregulated with respect to a reference if the average of Z-scores of that group is positive or negative, respectively.

## Supporting information

Metabolomics Data

Proteomics data

Sequences

## Acknowledgements

This research was supported by NIH grant 5R01GM068670 to E. I. Shakhnovich

## Author Contributions

Conceptualization, J.V.R and E.I.S, Methodology, J.V.R, and E.I.S, Investigation, J.V.R, Writing – Original Draft, J.V.R and E.I.S, Funding Acquisition, E.I.S

## Supplementary Information Text

### Supplementary experimental procedures

#### Construction of Plasmid pKD13-kefC(897-1863)-cmR-folA(wt)-kanR-apaH-apaG

A portion of chromosomal *apaH:apaG* genes was amplified by PCR using primers P4_chrom_flanking_for and PCRseq_apaH_rev. Then the pKD13 plasmid was linearized by PCR using the primers P4_chrom_flanking_for and PCRseq_apaH_rev. Both PCR products were purified and combined in a new PCR reaction using phosphorylated primers pKD13_post_downstream_for and PCRseq_apaH_rev. The linear product was ligated using T4 ligase to make plasmid Plasmid pKD13-*cmR*-folA(wt)-*kanR-apaH-apaG*. Then a portion of kefC gene was amplified by PCR using primers PCRseq_KefC_for2 and CapR-Chrom-Flanking rev. Then the plasmid pKD13-*cmR*-folA(wt)-*kanR-apaH-apaG* was linearized by PCR using primers Upstream_capR_for and pKD13_post_upstream_rev. Both products were combined in a new PCR reaction using phosphorylated primers PCRseq_KefC_for2 and pKD13_post_upstream_rev. Finally, the linear product was ligated using T4 ligase to make pKD13-*kefC*(897-1863)-*cmR*-folA(wt)-*kanR-apaH-apaG*

#### Mutations in D27 folA locus

Plasmid pKD13-*kefC*(897-1863)-*cmR*-folA(wt)-*kanR-apaH-apaG* was linearized using phosphorylated mutation primers D27Xmut_For and primer D27_rev.

#### Protein purification and characterization

D27 mutant fused to C-terminal (6x) His-tag were overexpressed using pFLAG expression vector under isopropyl β-D-1-thiogalactopyranoside (IPTG) inducible T7 promoter. The recombinant proteins were purified from lysates on Ni-NTA columns (Qiagen) as described previously (1).

#### Determination of kinetic parameters

DHFR kinetic parameters were derived from the analysis of progress-curve kinetics of NADPH oxidation in the presence of different concentrations dihydrofolate using the software Kintek Explorer (2) as described before (1). The data corresponds to the mean ± S.E. of at least three measurements.

#### bis-ANS Fluorescence Measurements

DHFR protein solutions (2 μM) in the presence of 12 μM of bis-ANS were prepared in 50 mM phosphate and 1 mM DTT at pH 7.0 and placed in a 1-cm path-length quartz cuvette. The samples were equilibrated for 5 min at 37 °C and the fluorescence emission spectra between 460 and 600 nm were recorded upon excitation at 395 nm. The emission band was integrated and the background of bis-ANS fluorescence in the absence of protein was subtracted. The area of the band was normalized to wild type E. coli. DHFR. The data corresponds to the mean ± S.E. of three measurements.

#### Determination of the melting temperature

DHFR solutions (5 μM) were prepared in 50 mM phosphate buffer and 1 mM DTT at pH 7.0 in the presence of 100 μM NADPH. A temperature ramp of 1 °C/min was set between 25 and 90 °C, and the tryptophan fluorescence was recorded by measuring the intensity at 370 and 320 nm upon excitation at 280 nm. Thermal denaturation curves were also performed in the presence of 5x concentration of Sypro Orange and 20 μM. Melting temperature and confidence intervals at 95% were determined from the simultaneous analysis of the thermal denaturation curves obtained with both tryptophan and Sypro Orange fluorescence using the online software Calfitter (3).

#### Measurements of total enzymatic activity in cell lysates

Cells from 14mL cultures grown at 30 °C at an OD of 0.5 were pelleted by centrifugation and lysed with 200μL of lysis buffer containing 1× Pop Culture reagent (Merck Millipore), 100 mM MES (2-(N-morpholino)ethanesulfonic acid) at pH 7.0 in the presence of 1 mM DTT in the presence of 1× complete protease inhibitor mixture (Roche) and 1 mg/mL lysozyme and incubated for 20 min at room temperature. The lysate was sonicated with 10 pulses of 1s,cleared by centrifugation and then 85μL of the soluble fraction was transferred to 96-well white plates for total activity measurements using a total assay volume of 100 μL. The lysate was preincubated with 100 μM NADPH and the reaction was started with the addition of 100 μM dihydrofolate and mixing. The decrease in fluorescence (excitation at 300 nm and emission at 440 nm) was measured over 60 min at 25 °C, and the initial velocities were computed. Measurements were also done in the presence of 1mM TMP to control for the non-DHFR-specific reaction. Data corresponds to the mean ± S.E. of three measurements.

### Predicting fitness of D27 mutants

The fitness of E. coli DHFR mutants can be predicted from in vitro biophysical parameters using a simple metabolic model described in an earlier work (1). The metabolic flux through DHFR reaction in DHFR mutant strains, normalized to wild type, can be approximated to:

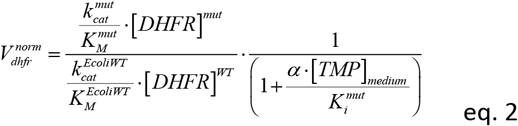

Where, 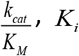 and_[*DHFR*]_ are, respectively, the catalytic efficiency, trimethoprim inhibition constant and intracellular protein abundance of the mutant or of the wild type, and [*TMP*]_*medium*_ and *α* are, respectively, the concentration of trimethoprim in the culture medium and the ratio of intracellular vs extracellular concentrations of trimethoprim, defined previously to be 0.1 (1). To connect flux to fitness, we first measure the growth rate of wild type *E. coli* cells at various concentrations of DHFR inhibitor trimethoprim.

Then fitness is plotted against the flux through DHFR reaction calculated at each concentration of inhibitor using equation 2, and catalytic parameters determined in vitro (Table 1). Protein abundance is determined by measuring the total catalytic activity in cell lysates, as described earlier (1). Finally, we use equation 3:

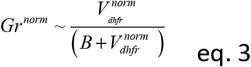

to fit the fitness vs V^norm^ data points to determine the parameter *B*, which represents the normalized flux at which growth rate is reduced by half. From this equation, the in vitro parameters determined for D27 mutants are used in equation 2 and 3 to predict fitness. The protein abundance of D27 mutants, however, cannot be determined from enzymatic assays in cell lysates, as these are inactive variants. Instead, protein abundance was predicted using an inverse relationship with relative bis-ANS fluorescence of those variants (1, 4). In qualitative terms, our model accurately predicted that the D27C variant is able to grow in the absence of folAmix supplement. However, the measured fitness was somewhat higher than the value predicted by the calculations. It is possible that some assumptions are not entirely met. One important assumption is that the concentration of DHFR and substrate dihydrofolate are either always constant or change only as a function of the flux calculated by equation S1. The latter implies that, for every given value of flux, the changes in DHFR and dihydrofolate concentration will always be the same, and thus always comparable regardless of what parameters are used as input in equation S1. However, this might not be always true. For example, a tight feedback control regulates the DHFR promoter activity, in response to the metabolic needs of the cell (5, 6). Likewise, a decrease in DHFR catalytic function is also expected to result in a considerable level of substrate build up. In these conditions, the magnitude of substrate *K*_M_ for each variant might become more relevant. In our calculations, however, substrate saturation is not considered. This may lead to deviations, especially when KM values differ significantly, as we find in this work. How much these dynamic processes may differently affect the flux through DHFR reaction in each mutant is still to be determined.

**Figure S1.**
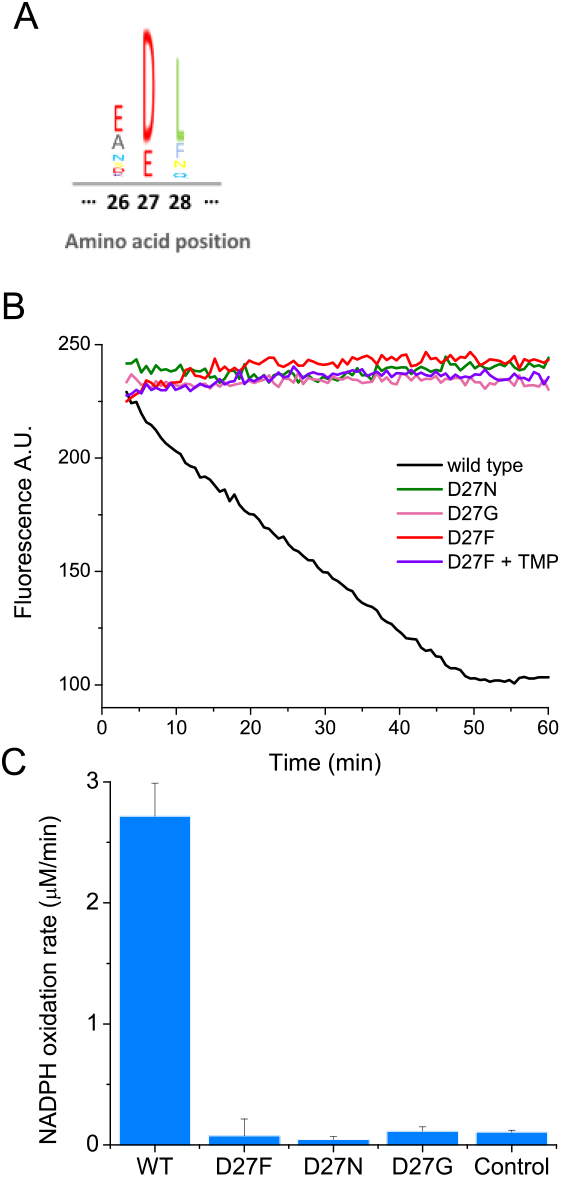
Mutations in key catalytic residue D27 impair DHFR function. A) Partial sequence alignment and sequence entropy for 290 bacterial DHFRs at the D27 locus showing strict conservation of carboxylate side chain residue. Sequence Logo for the alignment, generated using the MATLAB Bioinformatics Toolbox. B) D27 mutant strains lack measurable DHFR catalytic activity. Raw kinetic traces for the determination of DHFR catalytic activity in cell lysates from wild type and different D27 mutant strains and C) raw of NADPH oxidation rates measured from cell lysates from each strain. Cleared cell lysates were incubated with 100μM NADPH in 100 mM MES (2-(N-morpholino)ethanesulfonic acid) pH 7.0 in the presence of 1 mM DTT at 25°C and the reaction was initiated by the addition of 100 μM dihydrofolate. The decay in fluorescence intensity at 440 nm (excitation 300 nm) was measured over 1h, and the initial slopes were determined. Controls from non-DHFR-specific reaction were measured by the addition of trimethoprim (1 mM) to cell lysates.

**Figure S2.**
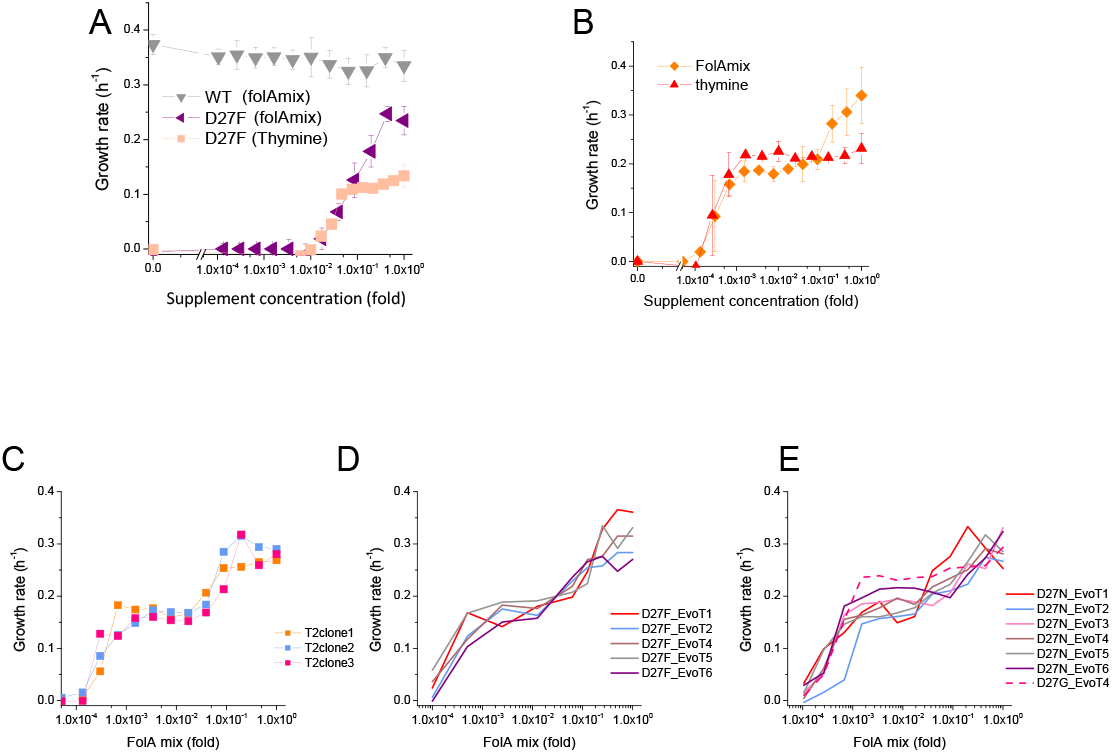
Naïve and evolved D27F require thymine for growth. A) Growth rate of naïve D27F in the presence of various concentrations of thymine, represented as fold dilutions with respect to initial concentration (200 μg/mL). For comparison, the growth profile in the presence of folAmix (at equivalent thymine concentrations) is represented for wild type and D27F mutant strains. Cultures were grown in M9-minimal medium at 30°C, and absorbance was monitored at 600nm. B) Evolved D27F strain (trajectory 1, cloneF51T1#1) was grown at different concentrations of either thymine or folAmix. C) Individual clones from the same evolutionary trajectory show similar fitness profile. Three randomly picked colonies from evolved strain D27F, trajectory 2, were grown at various concentrations of folAmix. D). Individual clones randomly picked form evolved cultures of different trajectories in D27F evolution show similar folAmix-dependent growth profile. E) Individual clones of evolved cultures from different trajectories in D27N and trajectory 4 in D27G evolution show similar folAmix-dependent growth profile as evolved D27F strains (panel D).

**Figure S3.**
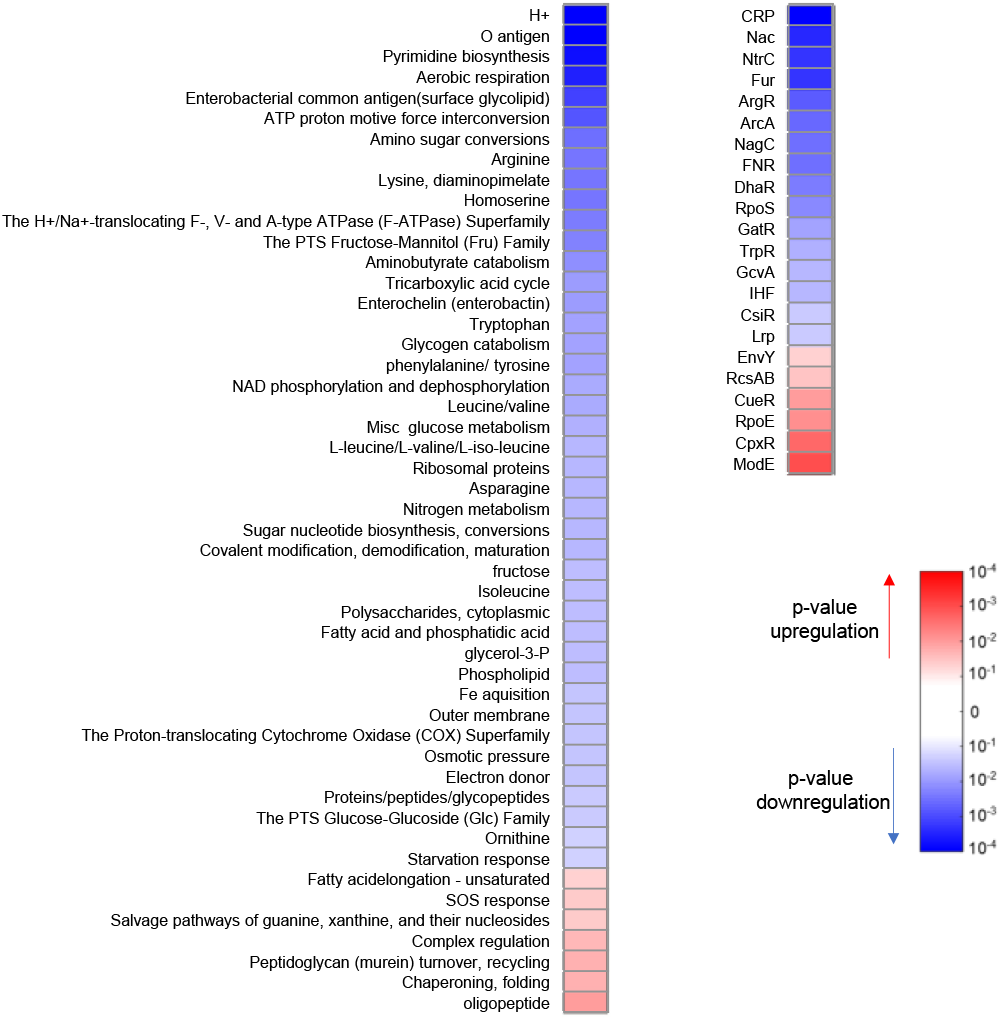
D27F mutation causes major proteomic changes. Variation in the abundance of proteins in naïve D27F strain, with respect to wild type, classified by functional and regulatory groups. For every group class (7), p-values of one-sample t-test were computed from the Z-scores of genes belonging to that category. A given group of proteins is considered to be upregulated if the average of the Z-scores is positive or downregulated otherwise. Only groups with significant changes (p<0.05) are represented.

**Figure S4.**
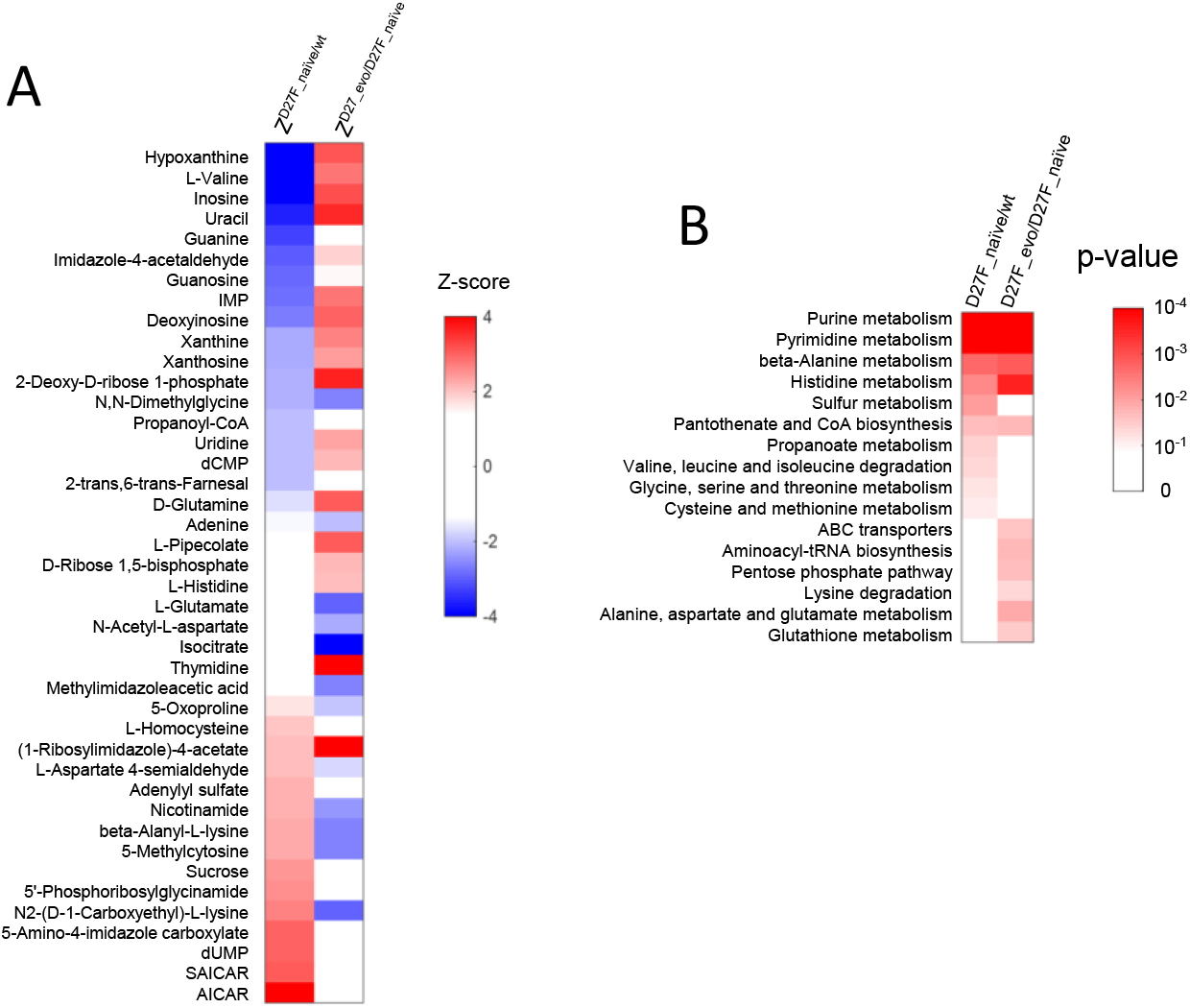
The abundance of various metabolites is strongly affected in naïve and evolved D27F strains. A) List of metabolites with highest absolute Z-scores in naïve and evolved D27F with respect to wild type and naïve D27F, respectively. B) Metabolic pathways associated with the most significant changes in naïve and evolved D27F, with respect to naïve D27F and wild type, respectively. P-values associated with each metabolic pathway were determined using a functional enrichment analysis (MBrole 2.0 online tool (8)) which compares, for every given class, the number of experimental hits and total background metabolites in that set. The metabolites with the highest absolute Z-scores (>1.96) were selected for the analysis.

**Figure S5.**
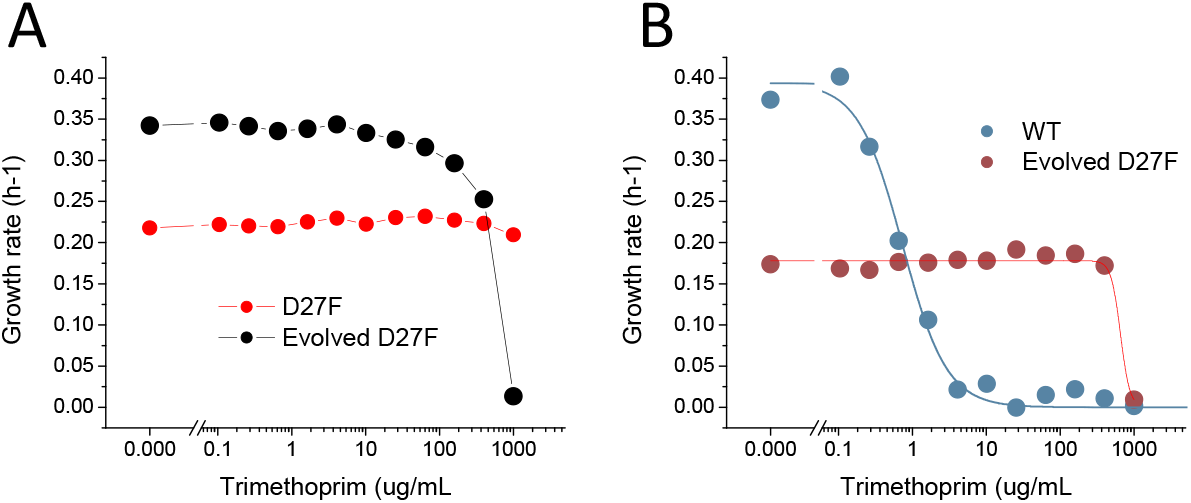
D27 mutant strains are extremely resistant to trimethoprim inhibition. A) Comparison of trimethoprim inhibition profiles of pre-evolved and evolved D27F in the presence of 1x folAmix. B) Comparison of trimethoprim inhibition profiles of wild type and evolved D27F strain at low concentrations of folAmix (5×10^−4^ fold). Solid lines are fits of logistic equations from which an IC50 of 0.8±0.2 μg/mL was obtained for wild type and 675±77 μg/mL was obtained for evolved D27F.

**Figure S6.**
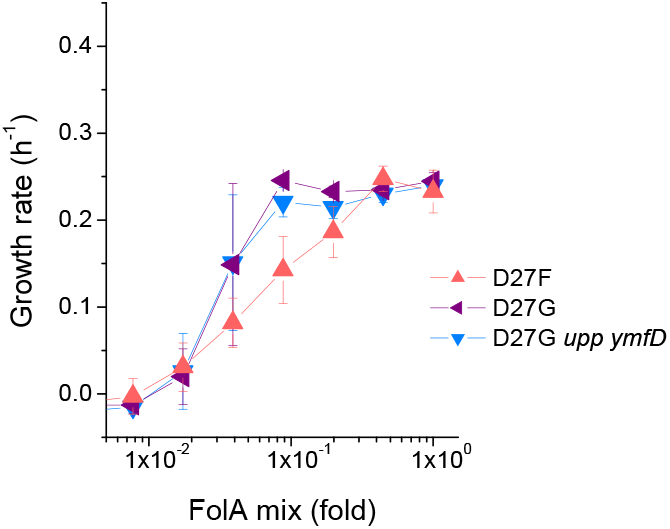
Mutations in *upp* and *ymfD* do not impact fitness of D27G. Strains where mutation D27G was introduced in folA gene were constructed and colonies were screened to select clones with intact *upp*, *ymfD* and *thyA* genes. The growth rate of 4 of these strains were measured at different concentrations of folAmix in M9-minimal medium at 30°C and compared with naïve D27G strain (*upp* p.17*STOP and ymfD p.A101V) used in the evolution study.

**Figure S7.**
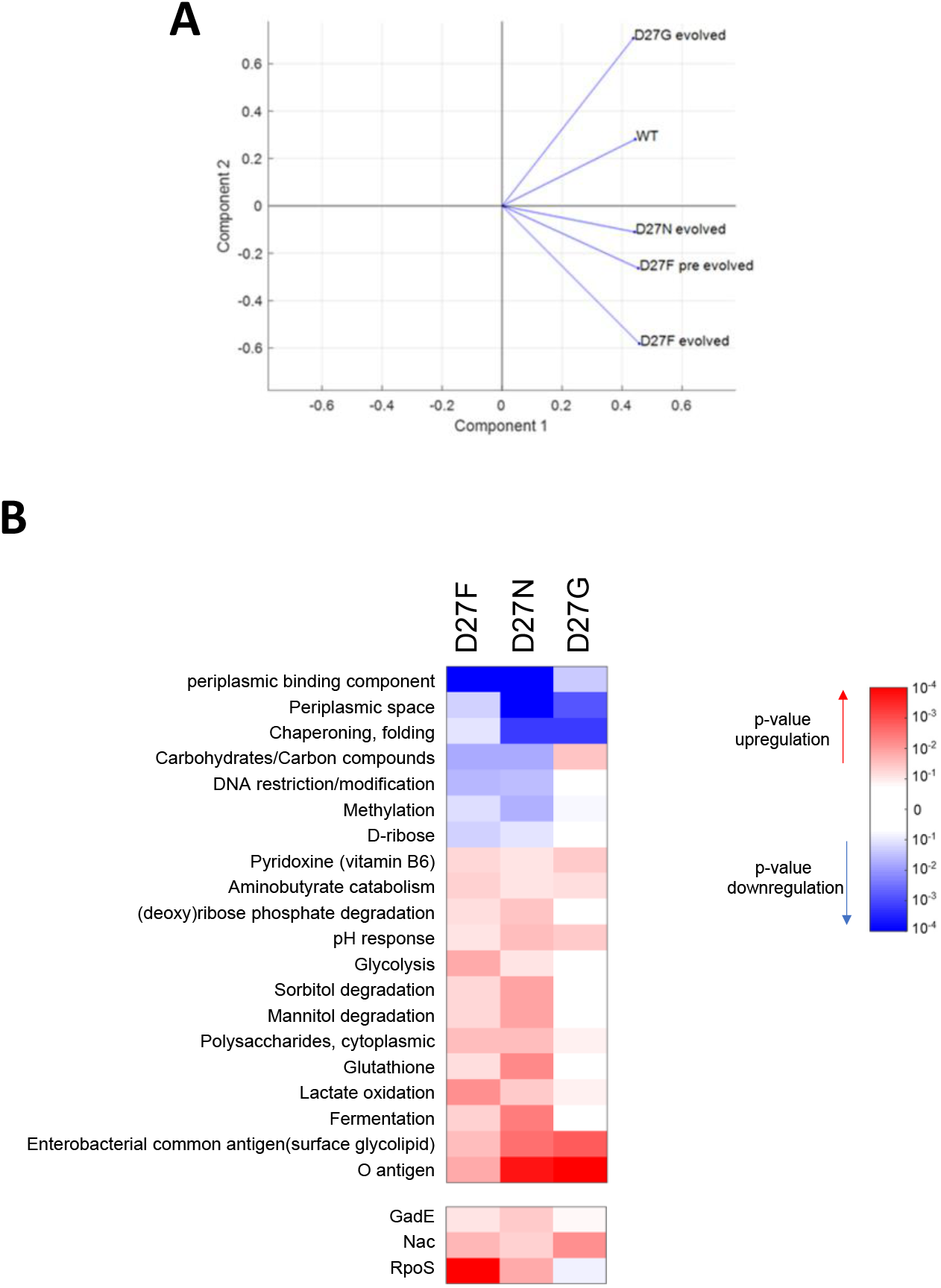
Proteomics profile of D27 mutants is associated with growth dependence on folAmix. A) PCA analysis of proteomics data shows that evolved D27G clusters close to wild type, in the space of two principal components, whereas folAmix-dependent strains D27 strain are found in a different quadrant. B) Variation in the abundance of proteins in evolved D27 strains, with respect to naïve D27F strain, grouped by protein functional and regulatory classification.

**Table S1–.**
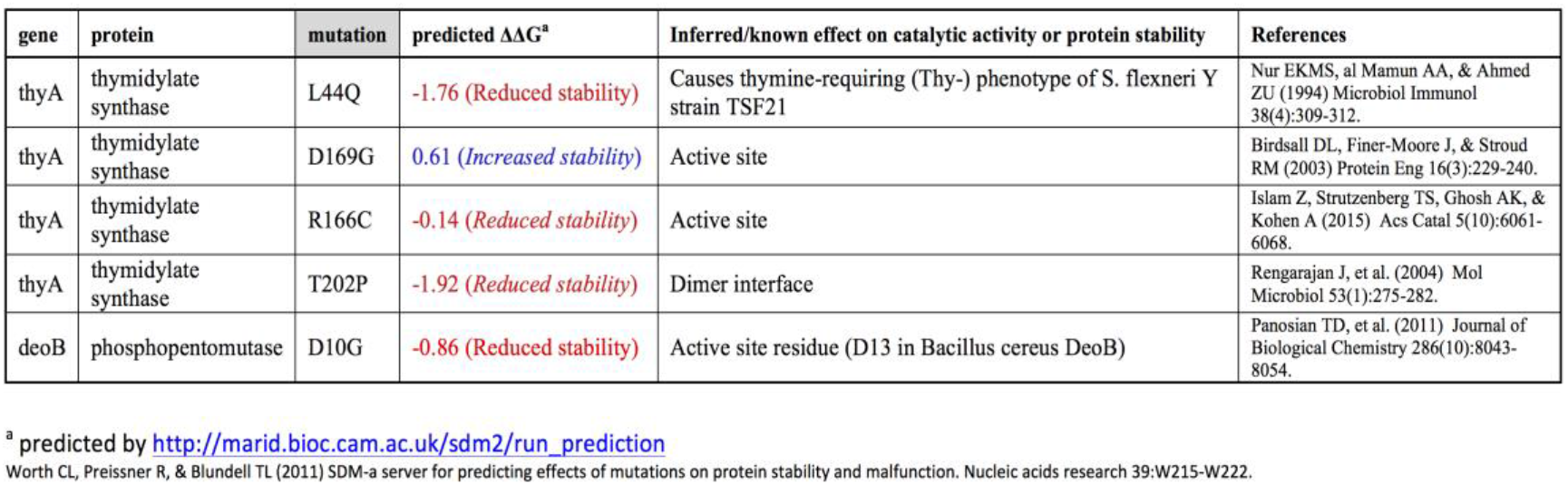
Single point mutations in thyA and deoB are predicted to impair catalytic activity and/or destabilize the protein

**Table S2–.** Resource table

| REAGENT or RESOURCE | SOURCE | IDENTIFIER |
| --- | --- | --- |
| Bacterial and Virus Strains |  |  |
| E coli Strain BW25113/pKD46 | E. coli Genetic Stock Center | strain#: 7739 |
| *E coli* Strain BW25113 ΔfolA::(*cmR-folA-kanR*) | This work | |
| *E coli* Strain BW25113 ΔfolA::(*cmR-folA.p.D27F-kanR*) | This work | |
| *E coli* Strain BW25113 ΔfolA::(*cmR-folA.p.D27N-kanR*) | This work | |
| *E coli* Strain BW25113 ΔfolA::(*cmR-folA.p.D27G-kanR*) | This work | |
| *E coli* Strain BW25113 ΔfolA::(*cmR-folA.p.D27C-kanR*) | This work | |
| TransforMax™ EC100D™ pir+ | Lucigen | Cat#ECP09500 |
| One Shot™ BL21 Star™ (DE3) Chemically Competent E. coli | Thermofisher | Cat#C601003 |
| Chemicals, Peptides, and Recombinant Proteins |  |  |
| Thymine | Sigma | Cat#T0376-10G |
| Inosine | MP biomedicals | Cat#102049 |
| Adenine | Sigma | Cat#A8626 |
| Methionine | Sigma | Cat#64313-25G-F |
| Glycine | Amresco | Cat#0167-1KG |
| Dihydrofolate | Sigma | Cat# D7006-25MG |
| NADPH | RPI | Cat# N20140-0.1 |
| Glucose | Fluka | Cat#49159-1KG |
| Difco M9 minimal salts, 5x | BD | Cat#248510 |
| Chloramphenicol | Calbiochem | Cat#220551 |
| Kanamycin sulphate | Teknova | Cat#K2105 |
| Recombinant protein E coli DHFR D27F | This work |  |
| Recombinant protein E coli DHFR D27N | This work |  |
| Recombinant protein E coli DHFR D27C | This work |  |
| Recombinant protein E coli DHFR D27G | This work |  |
| Critical Commercial Assays |  |  |
| KOD Hot Start DNA Polymerase | Sigma-Aldrich | Cat#71086 |
| Phusion High-Fidelity DNA Polymerase | Thermofisher scientific | Cat#F530S |
| DNA Clean & Concentrator (DCC) | Zymo research | Cat#D4013 |
| Oligonucleotides |  |  |
| Primers are listed in table S1 |  |  |
| Recombinant DNA |  |  |
| Plasmid pKD13 | E. coli Genetic Stock Center (9) | GenBank: AY048744.1 |
| Plasmid pKD13-cmR-folA(wt) | (10) |  |
| Plasmid pKD13-kefC(897-1863)-cmR-folA(wt)-kanR-apaH-apaG | This work |  |
| Plasmid pKD13-kefC(897-1863)-cmR-folA(p.D27F)-kanR-apaH-apaG | This work |  |
| Plasmid pKD13-*kefC*(897-1863)-*cmR*-*folA*(p.D27N)-*kanR*-*apaH*-*apaG* | This work |  |
| Plasmid pKD13-*kefC*(897-1863)-*cmR*-*folA*(p.D27G)-*kanR*-*apaH*-*apaG* | This work |  |
| Plasmid pKD13-*kefC*(897-1863)-*cmR*-*folA*(p.D27C)-*kanR*-*apaH*-*apaG* | This work |  |
| Plasmid pEM-Cas9HF1-recA56 | Addgene | Addgene plasmid # 89962 (11) |
| Plasmid pTRC-*thyA* | (12) |  |
| Plasmid pTRC-*tetR*-*deoB*(wt) | This work |  |
| Plasmid pTRC-*tetR*-*deoB*(p.309STOP) | This work |  |
| Plasmid pTRC-*tetR*-*thyA*(wt) | This work |  |
| Plasmid pTRC-*tetR*-*thyA*(755-762del) | This work |  |
| Software and Algorithms |  |  |
| Proteomics Grouping analysis, Matlab Code | (5) |  |
| Evolution Script, Freedom EVOware software | This work |  |
| Matlab R2018a | Mathworks | https://www.mathworks.com/products/matlab.html |
| Origin Pro7 | OriginLab | https://www.originlab.com/ |
| Freedom EVOware |  | https://lifesciences.tecan.com/software-freedom-evoware |
| Evolution Script, Freedom EVOware software | This work |  |
| MzMatch | (13) | http://mzmatch.sourceforge.net/installation.php |
| R 3.5.0 | The R Project for Statistical Computing | https://www.r-project.org/ |
| Ideom V19 | (14, 15) | http://mzmatch.sourceforge.net/ideom.php |
| Other |  |  |
| Freedom Evo 75 | Tecan | https://lifesciences.tecan.com/products/liquid_handling_and_automation/freedom_evo_series |
| Infinite® 200 PRO | Tecan | https://lifesciences.tecan.com/plate_readers/infinite_200_pro |
| Liconic STX44 | Liconic | https://www.liconic.com/index.php |

**Table S3.**
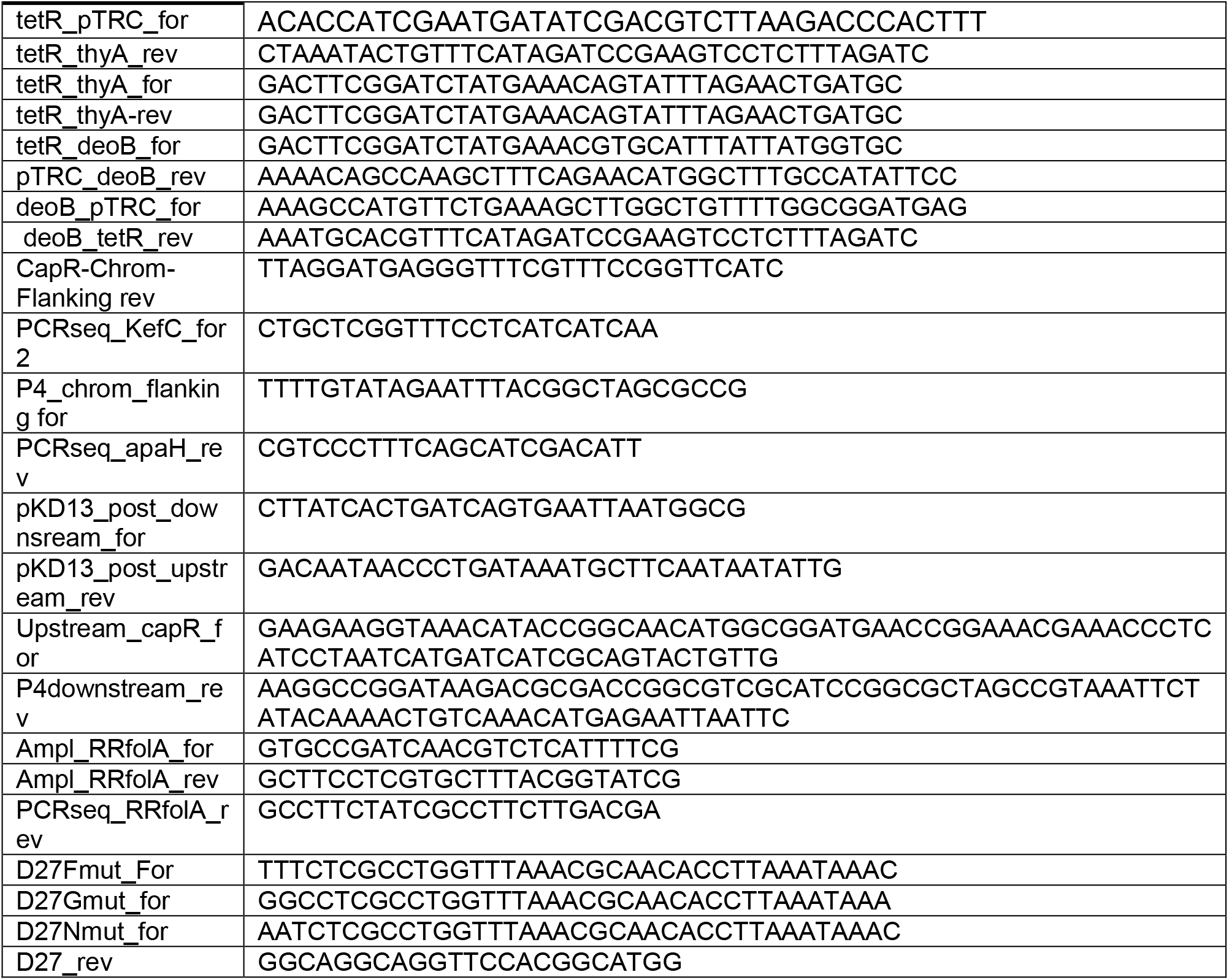
Primers list

**Dataset S1.** TMT proteomics of naïve and evolved D27 strains. All abundances are measured relative to WT strain which is taken as reference. Z-scores of abundance variation are calculated as ioutlined in text (see also (5))

**Dataset S2.** Variation of abundances of proteins grouped into functional groups according to (7). All protein abundances are relative to WT obtained from TMT proteomics (see dataset S1 for raw data). Average abundance variation z-score of proteins in the group and associated p-values are calculated as described in detail in (5). Groups marked in red represent functions that exhibit statistically significant increase or decrease of abundances in evolved but not naïve D27 strains.

**Dataset S3.** Variation of abundances of metabolites in naïve and evolved D27 strains.

**Dataset S4**. WGS and Sanger sequencing data for various evolutionary trajectories in naïve and evolved D27 strains.

